# Multivalent Anti-ACE2 Nanobodies Confer Broad Pan-Sarbecovirus Protection

**DOI:** 10.64898/2026.09.01.748598

**Authors:** Athanasios D. Bakasis, Rachel Patejak, Peter C. Fridy, Jesse Jenkins, Miranda Aldis, Viren A. Baharani, Lakshmi Sriram, Kelley R. Molloy, Brian T. Chait, Michael P. Rout, Frederick R. Cross, Paul D. Bieniasz, Theodora Hatziioannou

## Abstract

The continual emergence of SARS-CoV-2 variants that rapidly evade conventional spike-directed neutralizing antibodies, together with the ongoing risk of cross-species spillover and new sarbecovirus outbreaks, underscores the need to develop broadly acting, escape-resistant therapeutic agents. Here, we optimized a nanobody discovery pipeline incorporating competition-based yeast surface display assays to isolate single-chain variable heavy chain-only antibody domains (VHHs or nanobodies) that bind human ACE2 and inhibit SARS-CoV-2 entry. Dimeric VHHs, as well as bivalent and tetravalent Fc-fusion proteins exhibited markedly increased antiviral activity, blocking a broad panel of SARS-CoV-2 variants and diverse sarbecoviruses at low-nanomolar to picomolar concentrations. These agents did not affect ACE2 enzymatic function or cell surface expression. The VHH-Fc fusion proteins had favorable pharmacokinetics and conferred prophylactic protection in mouse models of both SARS-CoV-2 and SARS-CoV infection, showcasing their potential as broadly acting receptor-targeted biologics against pandemic-threat viruses.

## Introduction

Three coronaviruses have crossed into humans during the 21^st^ century, namely SARS-CoV, MERS-CoV and SARS-CoV-2 (1–3). Of these SARS-CoV-2 proved the most difficult to contain, resulting in the COVID-19 pandemic and over 7 million deaths world-wide (4). The wide distribution and diversity of sarbecoviruses circulating in non-human animals underscores the difficulty in predicting which sarbecovirus will emerge or cause disease in human populations, hampering preparedness efforts (5).

While the accelerated development of vaccines against SARS-CoV-2 was instrumental in eventually curtailing the pandemic (6, 7), the development of antiviral drugs as therapeutic or prophylactic agents has not advanced at the same pace. Neutralizing monoclonal antibodies (mAbs) against the spike protein of SARS-CoV-2 proved effective early in the pandemic but were derived from or closely related to those neutralizing antibodies commonly developed in humans following infection or vaccination(5, 8–10). Consequently, the continuous emergence of SARS-CoV-2 variants that evade naturally occurring neutralizing antibodies has rendered therapeutic neutralizing mAbs largely ineffective (11).

To overcome these limitations, we and others have developed mAbs that target the virus receptor, angiotensin-converting enzyme-2 (ACE2) which, as a host protein, is not subject to continuous and rapid evolution (12, 13). These mAbs bind human (hu)ACE2 with low-nanomolar affinities and block infection by diverse sarbecoviruses with potencies comparable to leading SARS-CoV-2 spike-directed therapeutic mAbs, but with far greater breadth (12). Resistance to such antibodies would require viruses to undergo major changes in their interaction with target cells, imposing a very high genetic barrier to escape. Moreover, receptors are frequently shared between members of virus families; indeed, huACE2 serves as a receptor for both SARS-CoV and SARS-CoV-2 and could do so for many animal sarbecoviruses that are potential pandemic threats (14–16).

An alternative avenue to conventional mAbs, are camelid single-chain variable heavy chain-only antibody domains, designated nanobodies or VHHs, that offer attractive properties for clinical applications (17, 18). VHHs are substantially smaller than full-length antibodies and are straightforward to engineer yet retain full antigen-binding capacity and, moreover, they have a specific ability to recognize buried or cryptic epitopes (19). Notably, VHH monomers can be covalently assembled into multivalent formats (e.g. dimers or fusions with a Fc domain of human immunoglobulin (IgG)) to enhance avidity, potency and improve pharmacology (20). They can also be fused to other agents to act as targeting moieties (21). Although VHHs targeting the SARS-CoV-2 spike protein exhibit potent antiviral activity, most are susceptible to spike mutations, sharing similar breadth limitations as traditional mAbs (22). Nevertheless, combining nanobodies creates synergy and at least partly alleviates mutation-dependent resistance.

Herein, we have developed pipelines for the identification and optimization of nanobodies targeting huACE2. Following immunization of llamas, huACE2-binding VHHs were identified using a mass spectrometry-based pipeline and an improved screening approach that employs competition-based yeast cell surface display (YD) assays to distinguish nanobodies that bind specific epitopes (23, 24). Candidate VHHs potently inhibited SARS-CoV-2 spike-mediated infection by targeting nonoverlapping epitopes within the huACE2-RBD interface. Optimization of VHHs via multimerization and fusion to immunoglobulin Fc domains increased antiviral activity, generating agents with both pan-sarbecovirus breadth and picomolar potency. Selected VHHs did not disrupt physiological huACE2 enzymatic activity or cell surface levels and conferred prophylactic protection against SARS-CoV-2 and SARS-CoV challenge in mouse models.

## Results

### Production and selection of VHHs targeting huACE2

To generate nanobodies against huACE2, a previously established pipeline combining mass spectrometry (MS) and high-throughput DNA sequencing was employed (24). Two llamas were subjected to four rounds of immunization at 21-day intervals with a recombinant soluble huACE2 extracellular domain (aa1–740), after which bone marrow plasma cells and sera were collected to generate a lymphocyte VHH cDNA library. Simultaneously huACE2-specific heavy chain-only antibodies were affinity-purified from sera, peptides generated by protease treatment and analyzed by liquid chromatography-tandem MS. The peptide sequences were matched to sequences from the VHH cDNA library to identify candidate huACE2-binding VHHs. Forty-nine candidate anti-huACE2 VHH sequences were expressed in bacteria and tested for binding to soluble recombinant huACE2 by ELISA (Fig. 1A). The majority (41/49) of anti-huACE2 VHHs bound huACE2 with EC50 equal or lower to 5nM. VHHs with the highest affinity were tested for antiviral activity against HIV-1 virus particles pseudotyped with SARS-CoV-2 spikes (Fig. 1B, Fig. S1A) (25). This assay identified one nanobody, NbA7, with strong antiviral activity, IC50 ∼12.8 nM, and three others, NbA15, NbA18, Nb19 with lower activity, IC50 ∼ 94.5, 94.1 and 88.8 nM, respectively (Fig. 1B).

**Fig. 1.**
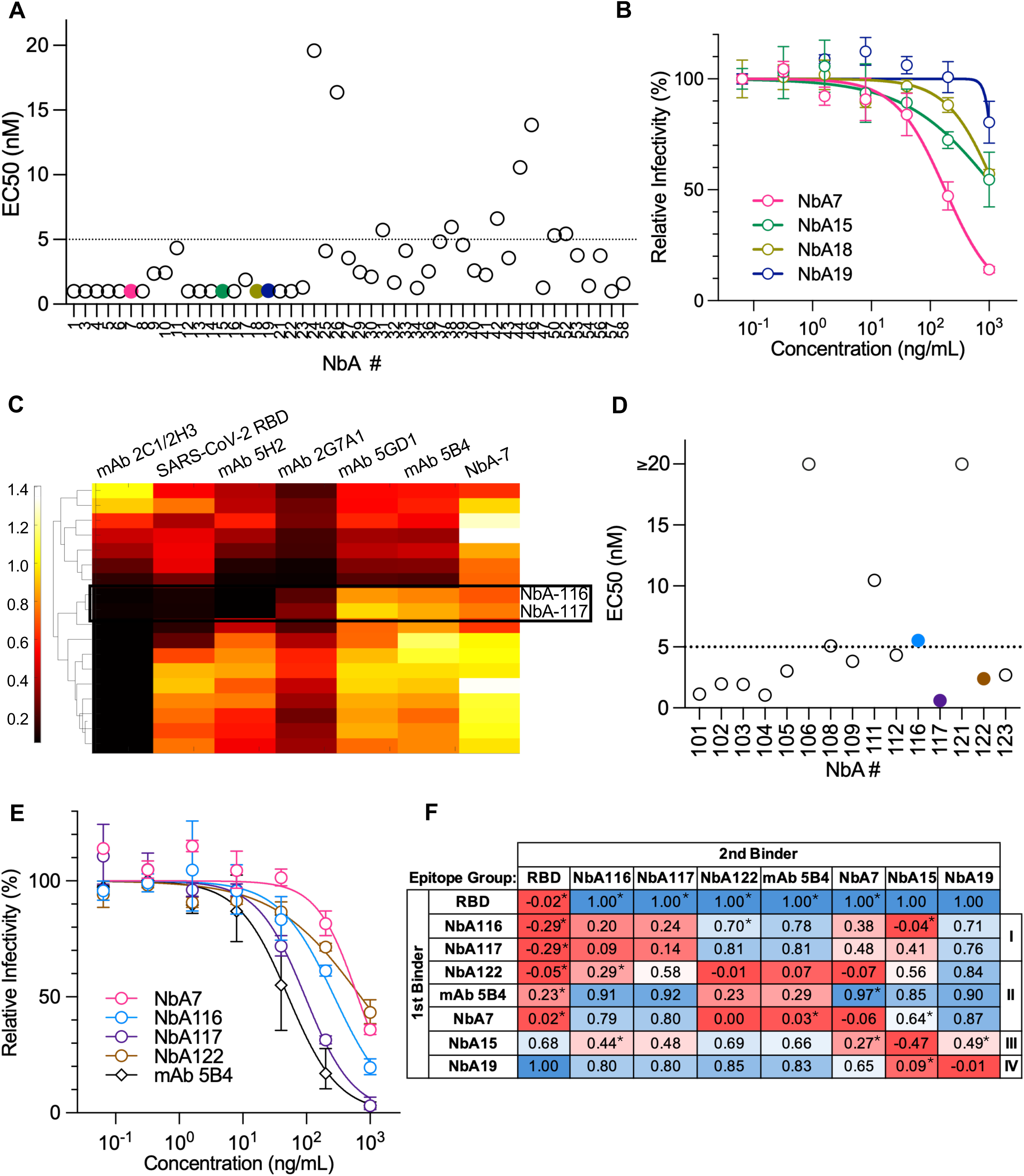
Discovery and characterization of anti-huACE2 VHHs with SARS-CoV-2 antiviral activity. **(A)** Binding of anti-huACE2 VHHs to huACE2. ELISA assays were performed with serial dilutions of 49 individual VHHs on microtiter plates previously coated with soluble recombinant huACE2. Detection of bound VHHs was performed with HRP-conjugated anti-VHH and EC50s were calculated by fitting a 4-parameter logistic curve. Selected VHHs with antiviral activity are colored as in (B). **(B)** VHH antiviral activity. Selected anti-huACE2 VHHs were serially diluted and incubated with Huh7.5 target cells and subsequently infected with an HIV-1 NanoLuc luciferase reporter virus pseudotyped with SARS-CoV-2 spike. Infectivity was determined by measuring luciferase activity at 48hrs post-inoculation. Values were normalized to infected, untreated positive control (set to 100%) and uninfected negative control (set to 0%). Data represent median and standard deviation from two independent experiments. **(C)** Yeast display competition experiments. Heatmap with hierarchical clustering (MATLAB clustergram, left) depicting the binding (read counts) of yeast cells expressing individual VHHs (rows) to huACE2-coated beads pre-saturated with the indicated known ACE2 binders (columns) normalized to binding to ACE2 in the absence of competitor. Color gradient reflects the level of competition; black (score 0) indicates complete block of binding by competitor, indicating an overlapping epitope; yellow-white (>=1) indicates no competition and distinct epitopes. **(D)** Binding of anti-huACE2 VHHs to huACE2. ELISA assays were performed as in (A) using VHHs identified via yeast surface display for specific competition with SARS-CoV-2 RBD. **(E)** Antiviral activity of selected VHHs derived from yeast display competition experiments as in (B). **(F)** Epitope binning of anti-huACE2 VHHs. Competition assays between the indicated VHHs, SARS-CoV-2 RBD or anti-huACE2 mAb 5B4 for binding to immobilized recombinant huACE2 were performed by SPR (surface plasmon resonance). huACE2 was immobilized on an SPR sensor surface and incubated with excess of binder 1 (capture). Binder 2 (competitor) was then flowed over the surface and binding (measured in resonance units) relative to binding in the absence of binder 1 was used to determine the level of competition. Heatmap color gradient reflects the level of competition with dark red signifying no binding of binder 2 in the presence of binder 1 (0 RU) and hence competition for the same epitope and dark blue equivalent binding to that in the absence of binder 1 (1 RU); that is binder 1 and binder 2 do not compete for the same epitope. Asterisks (*) denote cases where blocking effect is non-reciprocal. Nanobody epitope groups (I-IV) were defined by competition phenotypes.

Despite strong binding to huACE2, the majority of VHHs tested had little or no antiviral activity. This is likely due to targeting of epitopes that do not overlap with the SARS-CoV-2 spike binding interface and hence cannot inhibit spike binding. To improve VHH selection and identify a larger number of VHHs that target the huACE2–RBD interface, we adapted a yeast-display screening approach by incorporating competition for binding to the RBD interface (Fig. 1C, Fig. S1-2) (12, 23). A yeast-display library was generated using VHH cDNA from booster-dose–immunized llamas and yeast cells expressing VHHs were enriched for binding to huACE2 by incubation with huACE2-conjugated magnetic beads. The enriched library then underwent a second round of incubation with huACE2-conjugated magnetic beads that were previously incubated with SARS-CoV-2 spike RBD, NbA7, or one of several anti-huACE2 mAbs with potent antiviral activity, namely 5B4, 5GD1, 2G7A1, 5H2, and 2C1/2H3 (12). Following bead isolation, VHHs recovered from each condition were sequenced and the frequency of reads compared between the presence of a competitor to those recovered in the absence of competitor. The reduction in read frequency in the presence of a competitor indicated that the proteins recognized the same epitope (black in Fig. 1C) and were thus more likely to have antiviral activity (Fig. S1B). For example, VHH sequences with values near zero in the RBD column identify nanobodies that target the huACE2–RBD interface (Fig. 1C). Notably, although hierarchical clustering was based solely on competition phenotypes and not VHH sequences, we observed frequent, tight clustering of multiple VHH ‘family members’ presumably derived from somatic hypermutation of the same progenitor. For example, NbA116 and NbA117 were 94% identical at the amino acid level.

We selected representative VHHs that preferentially (a) competed with RBD, (b) had overlapping epitopes with other potent antiviral anti-huACE2 mAbs, and/or (c) did not share epitopes with NbA7 so as to expand our VHH repertoire. Binding of 15 selected candidate VHHs to huACE2 was verified by ELISA (Fig. 1D). Of the VHH that bound huACE2, three exhibited robust inhibitory activity against SARS-CoV-2 pseudovirus infection (Fig. S2A). Specifically, NbA117, NbA116 and NbA122 had IC50s of approximately 6.8, 19.6 and 50.9 nM, respectively (Fig. 1E), with NbA117 having the highest antiviral activity of all VHHs tested.

To determine whether selected VHHs targeted overlapping epitopes, we used surface plasmon resonance (SPR) to perform epitope binning (Fig. 1F). Soluble ectodomain of huACE2 was immobilized on a sensor surface and incubated with excess of either an individual VHH or soluble SARS-CoV-2 RBD or the anti-huACE2 mAb 5B4 (Binder 1). The ability of binder 1 to block binding of a second protein, binder 2, to huACE2 was then measured. This approach segregated the VHHs and mAb 5B4 into five distinct groups. Groups I–III (NbA116, NbA117, NbA122, and NbA7) all blocked RBD binding to huACE2. Within these groups, Group I (NbA116, NbA117) did not inhibit 5B4 binding to huACE2, while Group II (NbA122) and Group III (NbA7) did (Fig. 1F). Groups IV–V (NbA15, NbA19) inhibited neither RBD nor 5B4. NbA15 inhibited binding of NbA19 and partially inhibited binding of NbA116 and NbA117, indicative of either partially overlapping epitopes or steric interference due to epitope proximity on the ACE2. These findings were consistent with both the yeast display competition experiments and the antiviral potency of each VHH. Within these broad groups, further variation likely reflects different binding interfaces or affinities. Interestingly, the ability to compete for binding was dependent on the order of protein addition. For example, all nanobodies tested could bind to huACE2 when added as the binder 2 to ACE2 prebound to RBD, even though several of them completely inhibited RBD binding when added as binder 1 (Fig. 1F, top row). This ‘one way’ competition could reflect differences in affinity and/or footprint between VHHs and RBD. Structural predictions were carried out for these VHHs in complex with huACE2 using the AlphaFold-based AlphaRED pipeline (26). A high-confidence structure prediction was obtained for the NbA7-huACE2 interface (Fig. S3). The modeling suggests an NbA7 binding interface spanning huACE2’s α1-α3 helices that overlaps with both RBD and mAb 5B4 (PDB accession code: 8E7M) binding sites, consistent with the epitope binning competition experiments.

## Dimerization of anti-huACE2 VHHs increases their antiviral activity

We next reasoned that dimerization of nanobodies might increase antiviral activity by improving huACE2 avidity or steric obstruction of RBD binding, or a combination thereof. Therefore, homodimeric and heterodimeric VHHs were generated by incorporating a flexible Glycine-Glycine-Serine (GGS) linker between two VHH coding sequences. Heterodimerization improved antiviral activity to a greater extent than did homodimerization in all cases (Fig. 2A-B, Fig. S4). The orientation of the VHHs within heterodimers also impacted their antiviral activity, as evidenced by the different viral inhibitory potencies of heterodimers NbA15-117, IC50=178pM, and NbA117-15, IC50=965 pM (NbA15 and NbA117 at the N-termini, respectively). Heterodimers NbA15-117 and NbA15-7 were the most potent with IC50s of 149 pM and 178 pM, respectively, and were higher than the most potent anti-huACE2 mAb, 5B4 (IC50=224 pM) (Fig. 2A, Fig. S4A). Interestingly, the antiviral activity of the most potent individual VHHs targeting the ACE2-RBD interface, NbA7 and NbA117, was enhanced by fusion to NbA15, a VHH with lower antiviral activity that targets a different but likely proximal epitope (Fig. 1E-F).

**Fig. 2.**
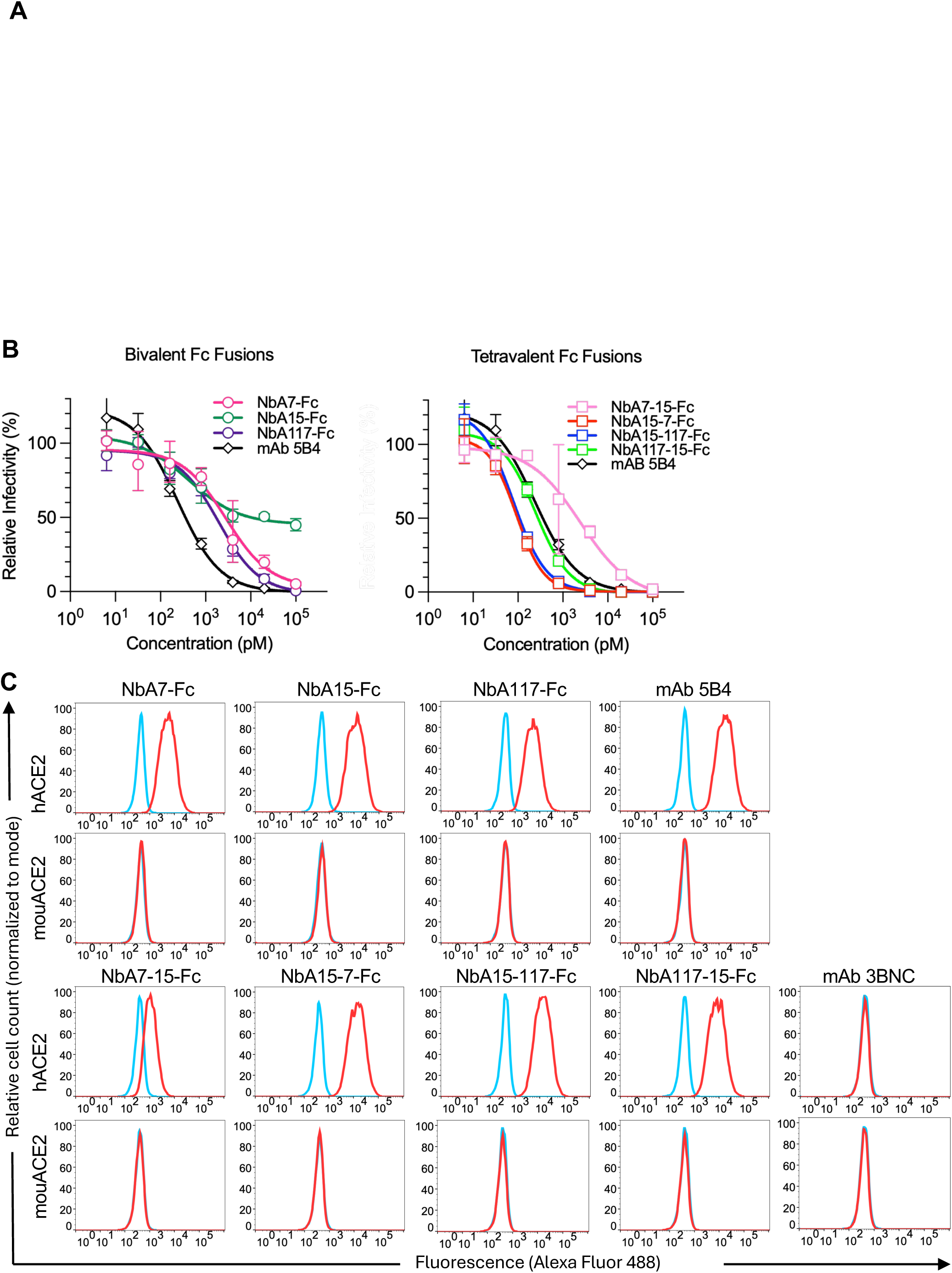
Dimerization of anti-huACE2 VHHs enhances their antiviral potency. **(A)** Antiviral activity of VHH homodimers and heterodimers. VHHs were dimerized by direct fusion of individual VHHs as indicated. Anti-huACE2 mAb 5B4 was included as control. Purified proteins were incubated with Huh7.5 cells and subsequently challenged with SARS-CoV-2-pseudotyped NanoLuc HIV-1 reporter virus. Infection was quantified 48hrs post-inoculation by measuring luciferase activity. Values were normalized to infected, untreated positive control (set to 100%) and uninfected negative control (set to 0%). Geometric mean and standard deviation are plotted. (**B)** Antiviral activity of VHH-Fc fusion proteins. Homodimeric and heterodimeric VHHs were fused to Fc domain of human IgG1. Infections were performed as in (A). **(C)** VHH-Fc binding to human and mouse ACE2. Parental A549 cells (blue) or A549 cells stably expressing huACE2, or mouACE2 (red) were incubated in the presence of the indicated anti-huACE2 VHH-Fc fusion proteins or mAb 5B4 (positive control) or anti-HIV-1 mAb 3BNC117 (negative control) at 0.5µg/mL. The cells were then incubated with Alexa Fluor 488 conjugated goat anti-human IgG and analyzed by flow cytometry.

We also generated bivalent and tetravalent VHH proteins by fusing monomer or dimer VHH proteins with the Fc domain of human immunoglobulin-γ1 (IgG1) (Fig. 2B, Fig. S4B). Tetravalent VHH-Fc fusion proteins had higher antiviral activity than all other proteins including Fc-fused and non-Fc VHH dimers and the previously identified anti-huACE2 mAb 5B4 (Fig. 2B, Fig. S4B). Specifically, IC50s values for the bivalent NbA7-Fc and NbA117-Fc were 2.9nM and 1.2nM respectively, while the tetravalent NbA15-7-Fc and NbA15-117-Fc had IC50 values of 70pM and 74pM, respectively. Tetravalent NbA15-7-Fc and NbA15-117-were also more potent than their corresponding non-Fc fused VHH heterodimers (IC50 = 137pM and 216pM for NbA15-7 and NbA15-117 respectively). The IC50s are comparable to or higher than those of potent spike-targeting conventional mAbs (11, 13, 27–30).

### Binding specificity of VHH-Fc fusion proteins

ACE2 proteins from multiple species can serve as receptors to SARS-CoV-2 with the notable exception of mouse (mou)ACE2, limiting the availability of mouse models for infection. We compared the ability of VHH-Fc fusion proteins to bind to the human cell line A549 engineered to express either huACE2 or mouACE2, at comparable cell-surface levels as measured by anti-HA tag staining (Fig. S5). Each VHH-Fc fusion protein bound A549 cells expressing huACE2 but not parental A549 cells nor those expressing mouACE2 (Fig. 2C). Tetravalent proteins NbA15-7-Fc and NbA15-117-Fc showed the highest level of binding to huACE2-expressing cells. Indeed, binding was nearly saturated at 0.1µg/mL (0.91 nM), consistent with their potent antiviral activity (Fig. S6).

### Anti-huACE2 VHHs-Fc proteins inhibit infection by multiple SARS-CoV-2 variants and diverse sarbecoviruses

Targeting the receptor rather than the viral spike is expected to increase the breadth of antiviral activity. Indeed, VHH-Fc proteins inhibited infection by all SARS-CoV-2 variants tested, including BA.2, BA.5, XBB.1.5, KP.2.3 (Fig. 3). In addition, they inhibited a cryptic wastewater variant termed ‘New Jersey Wastewater’ (NJWW) which is among the most highly divergent SARS-CoV-2 variants known, exhibiting only ∼80% amino acid identity with ancestral and delta linages and ∼77% with BA.2, BA.5, XBB.1.5 and KP.2.3 variants in the RBD (31). Importantly, inhibition potencies were comparable across variants (Fig. S4, Fig. S7A). Specifically, the IC50 values for NbA15-7-Fc and NbA15-117-Fc ranged from 60pm to 130pM and were similar to those against SARS-CoV-2 Wuhan strain (Fig. 2). NbA15-7-Fc and NbA15-117-Fc had comparable potency against the NJWW variant, despite its divergence, (IC50=105.8 and 113.5 pM, respectively), underscoring the resilience of receptor-targeting to spike sequence variation. Bivalent and tetravalent anti-huACE2 VHH-Fc–treated target cells were additionally challenged with viruses pseudotyped with spike proteins from SARS-CoV and from SARS-related coronaviruses from pangolin (CoV-GX), and bat (CoV-Rs4231, CoV-Rs7327). CoV-GX shares ∼92% and ∼75% amino acid identity with SARS-CoV-2, in the spike protein and RBD, respectively (32). CoV-Rs4231 S protein shows a ∼94% amino acid identity with SARS-CoV spike, sharing highly similarity in NTD but not RBD (33). All sarbecoviruses tested were inhibited by the human anti-huACE2 VHH-Fc fusions (Fig. S7B). As with SARS-CoV-2 variants, NbA15-7-Fc and NbA15-117-Fc exhibited the greatest antiviral potencies against divergent sarbecoviruses (IC50 = 25-141.2 and 37.3-250.8 pM, respectively) (Fig. 3B-D, Fig. S7B).

**Fig. 3.**
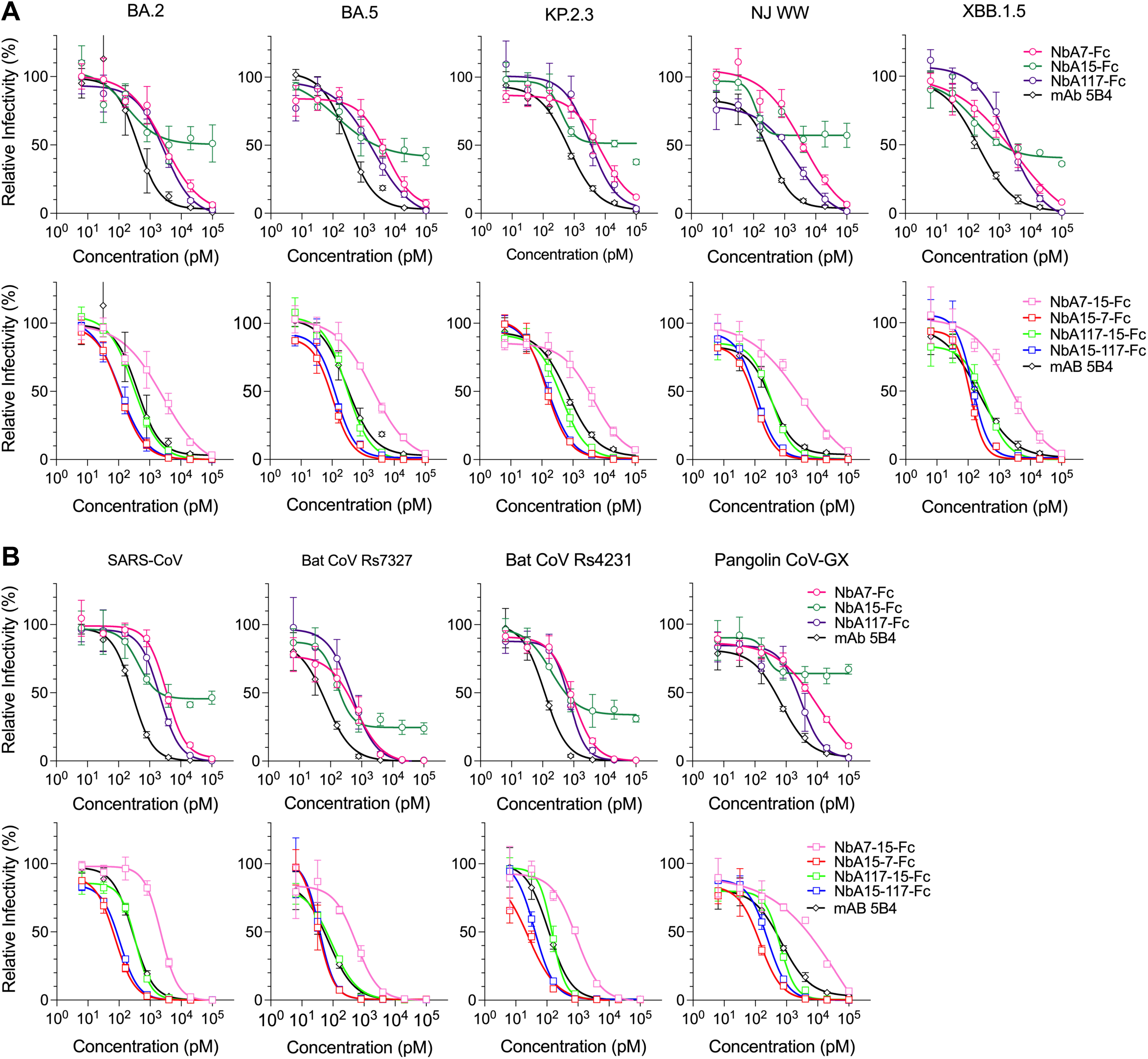
Anti-huACE2 VHH-Fc fusions block sarbecovirus spike pseudotyped infection. **(A)** VHH-Fc antiviral activity against SARS-CoV-2 variants. Bivalent and tetravalent VHH-Fc fusion proteins, harboring homodimers or heterodimers of NbA7, NbA15 and/or NbA117 were incubated with Huh7.5 target cells and subsequently challenged with NanoLuc HIV-1 reporter viruses pseudotyped with the SARS-CoV-2 variant spike proteins indicated. Infection was quantified 48hrs post-inoculation by measuring luciferase activity. Values were normalized to infected, untreated positive control (set to 100%) and uninfected negative control (set to 0%). Geometric mean and standard deviation are plotted. **(B)** VHH-Fc antiviral activity against divergent sarbecoviruses. As in (A) using NanoLuc HIV-1 reporter viruses pseudotyped with spike proteins from the divergent sarbecoviruses indicated.

### Genetic variation in huACE2 does not affect VHH binding

Non-synonymous variants of huACE2 at positions proximal to the SARS-CoV-2 spike-binding interface have been identified in humans at very low frequencies (0.001– 0.388%) (34). Such variation could, in principle, impact binding and antiviral activity of anti-huACE2 VHHs. We thus investigated whether the binding of VHH-Fc proteins to cells expressing huACE2 was affected by 18 naturally occurring amino acid substitutions located within the vicinity of the spike binding site (Fig. S8-9). Only 2 of the 18 substitutions, S19P and E329G, were predicted to potentially lower binding affinity to the SARS-CoV-2 spike (35). The majority of amino acid substitutions tested did not affect ACE2 recognition by NbA7-Fc, NbA15-Fc, NbA15-7-Fc, or NbA15-117-Fc. The exceptions were E35K and K68E, that reduced NbA117-Fc binding. However, heterodimerization of nbA117 with NbA15 and fusion to Fc restored binding to these huACE2 variants (Fig. S9). Of note, both substitutions are extremely rare in human populations (allele frequency = 1.64 x 10^-5^ and 1.09 x 10^-5^) (35). Consequently, naturally occurring ACE2 variation in humans is expected to exert minimal impact on the antiviral activity of the anti-huACE2 VHHs generated herein.

### Effects of anti-huACE2 VHHs on huACE2 enzymatic activity and cell surface expression

ACE2 catalyzes the hydrolysis of angiotensin II and its catalytic site is spatially separated from the N-terminal region that serves as the binding interface for sarbecovirus spike proteins, such that our nanobodies are not expected to interfere with ACE2 catalytic activity (36, 37). Nevertheless, to determine whether the anti-huACE2 VHH-Fc proteins affected huACE2 enzymatic function, we incubated Huh7.5 cells expressing huACE2 with each bivalent and tetravalent VHH-Fc protein, at the concentration range used in the viral inhibition assays, and measured huACE2 activity (Fig 4a). As expected, the small-molecule ACE2 inhibitor MLN-4760 inhibited enzymatic activity whereas the anti-huACE2 mAb 5B4 did not, consistent with our prior studies (12, 38). None of the anti-huACE2 VHH-Fc proteins tested inhibited huACE2 enzymatic function, even at the highest doses tested, corresponding to 100-1000xIC50 (Fig. 2). Therefore, VHH-Fc-bound huACE-2 fully retains enzymatic activity.

**Fig. 4.**
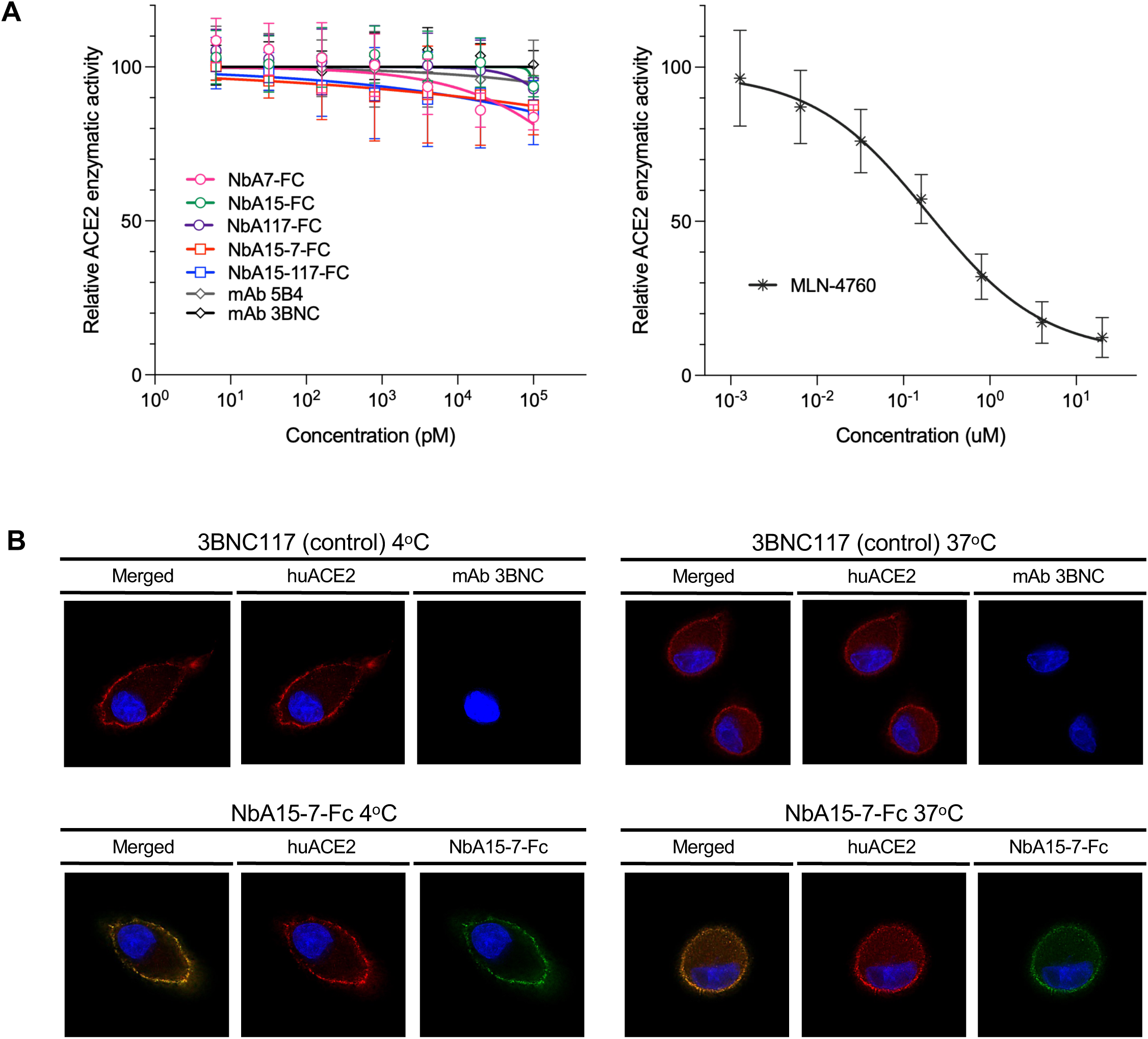
Effect of VHH-Fc fusion protein binding on huACE2 enzymatic activity and localization. **(A)** Effect of VHH-Fc binding to huACE2 enzymatic activity. Anti-huACE2 VHH-Fc proteins were serially diluted and incubated with Huh7.5 target cells that were subsequently treated with an ACE2 enzymatic activity indicator (left panel). Anti-huACE2 mAb 5B4 previously shown not to affect huACE2 enzymatic activity and a control anti-HIV-1 mAb 3BNC117 were included in the assays. An inhibitor of ACE2 enzymatic activity, MLN-4760, was used as positive control (right panel). ACE2 enzymatic activity was quantified by fluorogenic substrate cleavage and relative fluorescence units expressed as a percentage of the uninhibited (no-inhibitor) control, set to 100%. Mean and standard deviation are shown. **(B)** Effect of VHH-Fc on ACE2 cell surface localization. A549 cells expressing N-terminally HA-tagged huACE2 were incubated with VHH-Fc or a control anti-HIV-1 mAb 3BNC117 at 4°C or 37°C, as indicated. Staining of HA-huACE2 was performed with Alexa-594 (red) and VHH-Fc/mAbs with Alexa-488 (green). Cell nuclei were stained with DAPI (blue).

To determine whether VHH binding induced changes in huACE2 localization, we used a previously established fluorescence-based internalization assay (12). A549 cells expressing HA-tagged huACE2 were incubated for 1h, at either 4 °C or 37 °C, with Nb15-7-Fc or with a control anti-HIV-1 mAb 3BNC117 (39), then fixed and stained to visualize the cellular distribution of antibodies and the HA-tagged huACE2 (Fig. 4B, Fig. S10-12). As a positive control for internalization, cells were treated with an anti-CD44 antibody, known to undergo endocytosis (Fig. S10). At 4°C, no endocytosis was observed, HA-huACE2A was almost exclusively localized to the cell surface and colocalized with NbA15-7-Fc (Fig. 4B). At 37 °C, HA signals remained predominantly at the plasma membrane and also colocalized with NbA15-7-Fc. Importantly, intracellular HA-huACE2A localization and levels did not differ between NbA15-7-Fc–treated and control 3BNC117-treated cells (Fig. 4B, S10-12). These findings indicate that binding of the NbA15-7 moeity to huACE2 does not induce ACE2 internalization. This result minimizes the likelihood that such anti-huACE2 VHH-Fc proteins would undergo substantial target-mediated clearance from the circulation during *in vivo* use.

### Anti-huACE2 VHH-Fc proteins protect mice in SARS-CoV-2 and SARS-CoV models of infection

To evaluated VHH-Fc protein pharmacokinetics and prophylaxis *in vivo* we used three types of mice: 1) C57BL/6J (B6) mice that express only mouACE2, 2) huACE2 knock-in (huACE2-KI) mice in which huACE2 coding sequences have replaced the endogenous mouse *Ace2* locus and 3) huACE2-K18 transgenic mice that maintain mouACE2 expression but also express high levels of huACE2 under the control of the human keratin-18 (K18) promoter which allows expression in epithelia cells, including airway epithelia (40, 41). huACE2-K18 mice were crossed with IFNR−/− mice to generate huACE2-K18-IFNR^−/−^ transgenic mice.

To assess the pharmacokinetics of our most potent tetravalent VHH-Fc proteins, NbA15-7-Fc and NbA15-117-Fc, B6 and huACE2-KI mice were injected subcutaneously with 1.7 nmol of NbA15-7-Fc or NbA15-117-Fc (corresponding to the molar equivalent of a conventional mAb of ∼12.5 mg/kg). Serum samples were harvested on days 1, 4, and 7 post-injection and antiviral activity was measured using the SARS-CoV-2-pseudotyped HIV-1 reporter virus (Fig. 5A-B). Elimination rate constants (k) and half-life (t_½_) were calculated from the non-linear regression (one phase decay) analysis and for NbA15-7-Fc were k=0.09 day^-1^, t_1/2_=7.5 days in huACE2-KI mice and k=0.21 day^-1^, t_1/2_=3.3 days, in B6 mice. For NbA15-117-Fc they were k=0.23 day^-1^, t_1/2_=3 days in huACE2-KI and k=0.19 day^-1^, t_1/2_=3.6 days in B6 mice. NbA15-7-Fc demonstrated more favorable PK properties than Nb15-117-Fc and in mice injected with NbA15-7-Fc, serum concentrations remained well above 10µg/ml on day 7 and were comparable to typical human mAb levels in mice (42, 43). Neither of the VHH-Fc proteins produced observable adverse effects in the mouse strains tested.

**Fig. 5.**
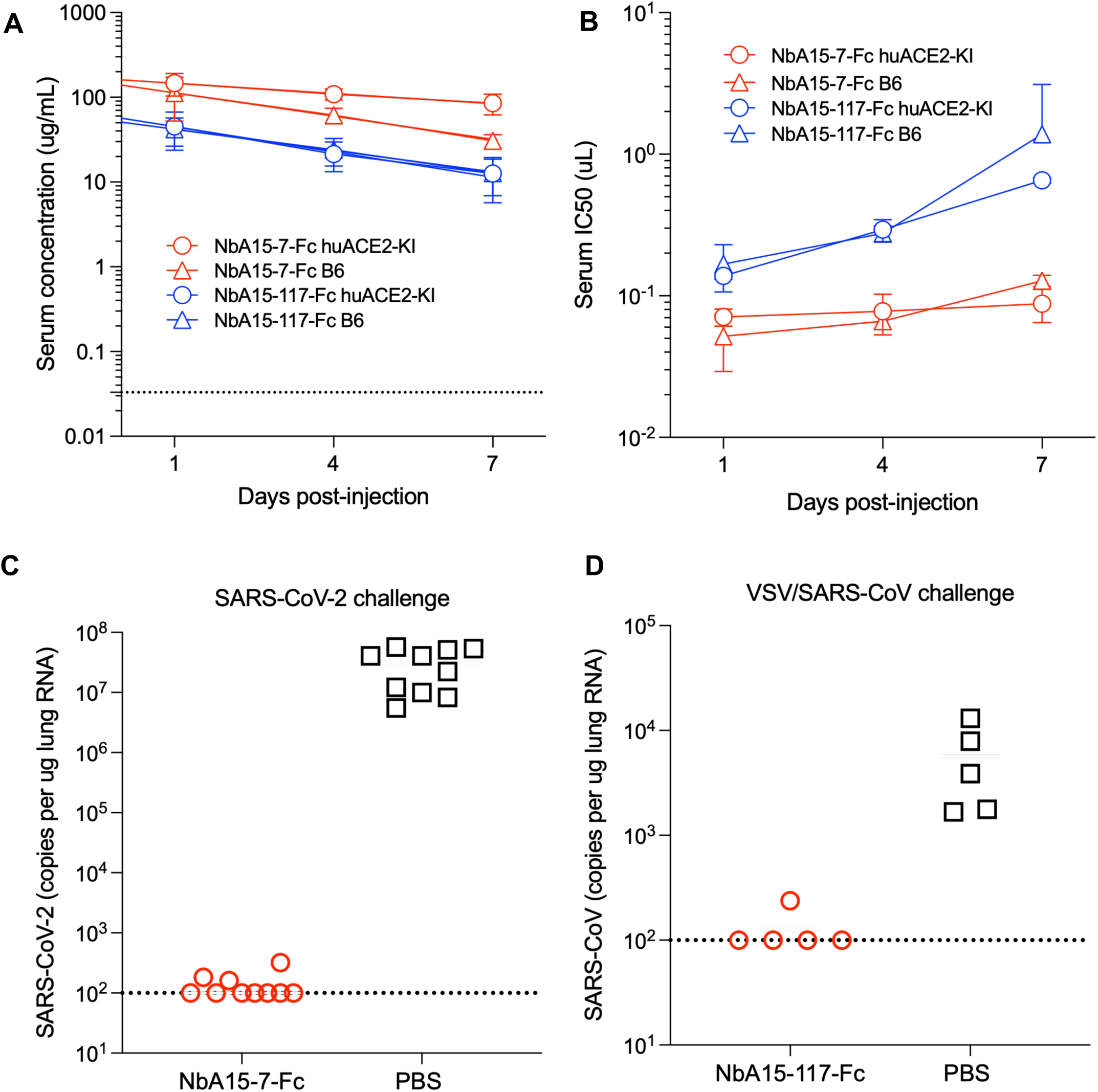
Anti-huACE2 VHH NbA15-7-Fc fusion confers protection against SARS-CoV-2 infection in mice. **(A)** Pharmacokinetics of anti-huACE2 VHH-Fc fusion proteins in mice. NbA15-7-Fc and NbA15-117-Fc were injected subcutaneously at 1.7 nmol/mouse (∼180 μg) into huACE2-KI and wild-type B6 mice (n=3 per group). Dilutions of sera collected on days 1, 4, and 7 were tested for viral inhibition activity, along with NbA15-7-Fc and NbA15-117-Fc protein dilutions as standards. Relative inhibition curves for sera and standards enabled extrapolation of mouse serum concentrations. Data presented as mean values and standard deviation for each group and timepoint. Lines represent non-linear regression analysis (one phase decay) for calculating elimination rate constants (k) and half-life (t_½_). The dashed line denotes the VHH-Fc fusion protein IC50s for pseudovirus inhibition in cell culture. **(B)** Serum antiviral activity. Serum volumes (in μl) required to inhibit 50% of the maximum virus infectivity in (A) were calculated for each mouse and each time point. Mean values and standard deviation for each group of mice over time are plotted. **(C)** HuACE2-KI mice were injected intraperitoneally with 1.7nmol/mouse (∼180 μg) of NbA15-7-Fc and 2 days after injection each mouse was challenged intranasally with 2 × 10^5^ PFU of SARS-CoV-2 (USA_WA/2020 P). Three days post-challenge mouse lungs were collected, RNA extracted, and the number of SARS-CoV-2 genomes per µg of total lung RNA was measured using qRT–PCR. Horizontal dashed line indicates the limit of detection (LOD) at 100copies/μg. **(D)** huACE2-K18-IFNR^−/−^ mice were injected intraperitoneally with 1.7 nmol/mouse (∼180 μg) of NbA15-7-Fc. Two days post-injection, each mouse challenged intranasally with 2 × 10^4^ PFU VSV/SARS-CoV chimeric virus (virus titers measured on Vero E6 cells). Two days post-challenge mouse lungs were collected, RNA extracted and the number of VSV viral genomes per µg of total lung RNA was measured using qRT–PCR. Horizontal dashed line indicates the LOD at 100 copies/μg.

To assess the ability of VHH-Fc to protect against SARS-CoV-2 infection, NbA15-7-Fc was administered intraperitoneally to huACE2-KI mice (n = 10 per group) at 1.7nmol/animal. Two days post-inoculation mice were challenged intranasally with 2×10^5^ PFU (plaque-forming units) of SARS-CoV-2 (strain USA_WA/2020) (Fig. 5C). Mice were bled immediately before challenge, and serum was analyzed to confirm adequate and comparable drug administration (Fig. S13). At 2 days post-infection, levels of viral RNA in the lungs of mock-treated mice ranged from 10^6^-10^8^ copies/μg total RNA. Administration of NbA15-7-Fc markedly decreased SARS-CoV-2 RNA levels in the lungs; viral loads were reduced to levels that were at or below the limit of detection, mirroring the efficacy reported for mAb 5B4 at an equimolar dose (Fig. 5C). These findings demonstrate that prophylactic use of anti-huACE2 VHH-Fc proteins in huACE2-KI mice confers near-sterilizing protection against high dose SARS-CoV-2 intranasal challenge.

In cell culture assays with pseudotyped virus particles, NbA15-7-Fc has broad antiviral activity against diverse sarbecoviruses including SARS-CoV (Fig. 3B). The latter virus requires handling under strict laboratory safety conditions that limit the ability to test interventions. To overcome this limitation, we developed a modified mouse challenge model based on a replication-competent chimeric VSV that expresses the SARS-CoV spike protein instead of the native glycoprotein G, designated VSV-SARS-CoV. Intranasal inoculation of huACE2-K18-IFNR^−/−^ with the chimeric VSV/SARS-CoV results in ∼10^4^ copies VSV RNA copies per μg of total lung RNA at 2 days post-infection, as previously reported (44). Mice were injected intraperitoneally (6 mice per group) with 1.7nmol of NbA15-7-Fc per animal, 2 days prior to intranasal challenge with 1×10^6^ PFU of VSV/SARS-CoV (Fig. 5D). Mice were bled immediately before challenge, and serum was analyzed to confirm adequate and comparable drug levels between mice (Fig. S13). In contrast to control mice, that had viral copies between 10^3^ to 10^4^, administration of NbA15-7-Fc resulted in viral loads below the limit of detection in the lungs of 5/6 mice. In one mouse viral loads were detectable but 10-fold lower than control mice (Fig. 5D). This data demonstrate that anti-huACE2 VHH-Fc fusion proteins confer protection against viruses with an entry pathway dependent on SARS-CoV spike.

## Discussion

Here we show that identifying VHHs that target the receptor-spike interface of huACE2 can provide the basis for building novel antiviral tools. Notably, generation of heterodimers of nanobodies that target distinct -but spatially close epitopes – resulted in the generation of fusion proteins with greatly improved antiviral activity. This activity was increased even further by the generation of tetravalent proteins by fusing VHH heterodimers to Fc. Such VHH–Fc proteins have a broad range, inhibiting infection by viruses bearing spikes from all SARS-CoV-2 variants and divergent sarbecoviruses tested. Additionally, VHH-Fc do not affect the enzymatic activity or cellular distribution of huACE2. Importantly, VHH-Fc with favorable pharmacokinetics in mice, provided almost sterilizing protection against a high-dose intranasal SARS-CoV-2 challenge. Furthermore, using our newly developed challenge model with a chimeric VSV expressing the SARS-CoV spike, we demonstrate that VHH-Fc fusion candidates also protect against a virus that mimics SARS-CoV entry. These properties render ACE2-targeting biologics attractive candidates for both prophylactic and therapeutic applications.

Using a mass spectroscopy-guided VHH discovery pipeline together with a refined yeast-display assay that employs competitive screening, this study introduces a platform that selectively enriches VHHs targeting the receptor-ligand interface from an excess of “irrelevant” non-blocking epitopes (23, 24). Compared with conventional screening methods that require testing of individual candidates, this design is advantageous and permits simultaneous identification of a repertoire of VHHs with non-overlapping, ligand-blocking specificities that could later be deployed as cocktails or in multi-specific formats. This approach mirrors the design of caplacizumab, an approved VHH-Fc-based therapy used to treat acquired thrombotic thrombocytopenic purpura (20). In doing so, it combines the advantages of both approaches: nanobodies that are intrinsically potent and synergize when assembled into multivalent formats, and an Fc domain that confers *in vivo* advantages such as extended half-life, improved pharmacokinetics, and compatibility with standard antibody manufacturing and commercialization platforms (17, 18). Our study provides mechanistic insights that increasing valency markedly amplifies antiviral activity without compromising ACE2 function. Dimerization of VHHs that target sterically adjacent epitopes substantially improves antiviral potency, likely due to a combination of avidity effects and more effective steric occupancy of huACE2 at the spike interface. Conversion to bivalent, and particularly to tetravalent, VHH-Fc fusions further enhances potency to levels comparable to or exceeding those of anti-spike and anti-huACE2 mAbs. Observing that orientation and pairing of VHHs within heterodimers influence antiviral potency underscores the importance of geometry in tertiary and quaternary structural space for protein–protein interactions.

Anti-ACE2 mAbs, established previously by our lab and others, demonstrate that blocking the spike-ACE2 interaction could provide broad protection across diverse sarbecoviruses by creating a high genetic barrier to viral escape (12, 13, 45). Indeed, this pipeline has led to the identification and generation of nanobody fusion proteins with very high antiviral activity that improve on a prior report generating anti-ACE2 nanobodies with antiviral activity (45). Direct comparison of the potency of our best nanobody candidates with a recently described anti-ACE2 nanobody, B07-Fc, is difficult as antiviral activity against virus infection was not measured directly but instead evaluated based on cell-to-cell fusion assays. Furthermore, B07-Fc provided protection against intranasal SARS-CoV-2 challenge in less than 50% of the mice tested. In contrast, our best candidate, NbA15-7-Fc described in our study protection against intranasal challenge with SARS-CoV-2 in 100% off animals and against a surrogate model for SARS-CoV infection in 80% of the animals (Fig. 5C,D).

From an evolutionary perspective, directing inhibitors to ACE2 rather than to spike is predicted to impose a substantially higher barrier to viral escape. The VHH-Fc fusions inhibit a broad panel of SARS-CoV-2 variants, including highly antigenically drifted lineages, as well as SARS-CoV and several animal pandemic-threat sarbecoviruses. Potency is similar across a diverse set of sarbecoviruses and the limited impact of huACE2 polymorphisms near the receptor–spike interface suggests that naturally occurring huACE2 variation is unlikely to dampen efficacy in human populations. Furthermore, while VHH-Fc fusions are well-suited for systemic administration as injectables, the smaller VHH dimers, by virtue of their superior tissue penetrance and low molecular weight, represent compelling candidates for intranasal or inhaled delivery. This could have the advantage of providing anti-viral protection at the primary site of virus entry. Finally, the approaches developed herein could be applied to other viral receptor proteins, providing a readily adaptable platform for the development of receptor-targeted prophylactics and therapeutics.

## Materials and Methods

### Cell lines

Human embryonic kidney HEK-293T cells (American Type Culture Collection (ATCC), CRL-3216), human hepatoma-derived Huh-7.5 cells (46), and A549 cells (adenocarcinomic human alveolar basal epithelial cells) were maintained in Dulbecco’s modified Eagle medium (DMEM) supplemented with 10% fetal bovine serum (Sigma, F8067) and gentamicin (Gibco) at 37 °C and 5% CO₂. A549 cells stably expressing human huACE2 or mouACE2, each bearing a C-terminal 3×HA tag, were previously generated using the retroviral vector pLBCX and selected with blasticidin, as described (13). A549 cells stably expressing human huACE2, bearing a N-terminal 1×HA tag, were previously generated and described in (12). Suspension-adapted HEK-293T cells (Expi293T; Thermo Fisher Scientific; A40003643) were maintained in serum-free Expi293 Expression Medium (Thermo Fisher Scientific) in vented flasks at 37°C and 8% CO₂ with orbital shaking. All cell lines were monitored periodically for the absence of mycoplasma and adventitious retroviral contamination.

### Generation and identification of huACE2-binding VHHs

To generate huACE2-targeting VHHs, two llamas were immunized subcutaneously four times at 21-day-intervals with recombinant huACE2 extracellular domain (residues 1–740) fused to a polyhistidine tag (300 µg per animal), as previously described (12, 23, 24). Following immunization, nanobody identification was performed according to an established mass spectrometry-based protocol (24, 47). Briefly, BMPCs and sera were collected to generate an *in silico* VHH library by sequencing VHH domains PCR-amplified from lymphocyte-derived cDNA with Illumina MiSeq 2×300bp pair-end reads. HuACE2-specific HCAbs containing the VHHs of interest were affinity-purified from sera. Peptides produced by trypsin and chymotrypsin digestion were analyzed by liquid chromatography–tandem mass spectrometry (LC-MS/MS) and matched to library sequences to identify candidate huACE2-binding VHHs.

To broaden and diversify the VHH repertoire, the llamas underwent four additional boost immunizations with 400 µg recombinant huACE2. BMPCs were collected again to generate VHH cDNA libraries, employing a modified yeast-display selection method (23). Yeast display library generation was performed as described, with VHH PCR-amplified from cDNA using oligos providing flanking homology to the yeast display vector, for subsequent gap-repair and yeast transformation (48, 49). For surface nanobody induction, 1 ml of stationary-phase yeast culture was diluted into 9 ml of 6% galactose– 0.2% galactose yeast minimal medium (ScMin), grown at 30 °C for 20–24 hours, supplemented with 10% fresh minimal medium and 3% galactose, and incubated at 30 °C overnight. These conditions were sufficient for nanobody expression, as demonstrated by binding purified GFP to a concurrently induced anti-GFP yeast-display control and measuring fluorescence. A population of cells in these anti-GFP controls did not display fluorescence; prior experiments suggested that these cells had experienced plasmid loss events under galactose induction. Because these cells lacked plasmids, they did not express nanobodies and therefore did not contribute to downstream processes; their presence during ACE2-binding nanobody capture was inconsequential. For yeast affinity capture, purified huACE2 was conjugated to M-270 Carboxylic acid Dynabeads (Thermo Fisher Scientific) using EDC/NHS coupling according to the manufacturer’s protocol, with minor adaptations; the binding and washing conditions were detailed previously (23). Details of the optimized binding procedure are as follows: 1 ml of induced yeast library was pelleted and resuspended in BPT buffer—composed of 1% BSA (Fraction V, protease-free; GoldBio, St Louis, MO), 1× PBS (137 mM NaCl, 2.7 mM KCl, 10 mM Na2HPO4, 1.8 mM KH2PO4, pH 7.4), and 1% Tween-20 (Sigma Aldrich). Cells were incubated with 1.45 mg of huACE2-conjugated Dynabeads and rotated at 30 °C for 1 hour. This was followed by five washes in BPT buffer of the bead-bound cells on a Dynal MPC-6 magnetic stand, with samples kept at 2 cm from the magnet and a 5 min binding per wash. For the epitope-binning experiments, affinity capture of the yeast-display library was performed as described above, except that each 1 mg aliquot of huACE2-conjugated Dynabeads was prebound by adding 40 μg RBD or an anti-ACE2 nanobody, or 200 μg of an anti-huACE2 monoclonal antibody in BPT buffer and rotating for 1 hour at room temperature. Affinity capture with ACE2-conjugated Dynabeads and subsequent nanobody isolation were performed as normal. For sequencing of nanobody clones in the purified yeast library, after binding, the beads with bound cells were transferred to ScMin– 2% glucose and incubated for 30–48 h. Cells were pelleted and lysed with Zymolyase; DNA was then purified on Qiagen miniprep columns following the manufacturer’s protocols. The DNA preparation was amplified with sequencing primers and sequenced at the Rockefeller Genomics Facility on an Illumina NextSeq 2000 (P1: 2 × 300bp reads).

### Human ACE2 and sarbecovirus spike expression plasmids

Plasmids encoding the spike proteins of SARS-CoV; SARS-CoV-2 Wuhan-Hu-1 (bearing the furin cleavage-site mutation R683G); clinically relevant Omicron subvariants BA.2, BA.5, XBB and KP.2.3; and the pangolin (*Manis javanica*) coronavirus from Guangxi, China (pCoV-GX) and rufous horseshoe bat (*Rhinolophus sinicus*) coronaviruses Rs4231 and Rs7327 were cloned into a pCR3.1 backbone and have been previously described (12). The plasmid encoding the New Jersey wastewater–derived SARS-CoV-2 spike, cloned into a pCR3.1 expression vector, is described in (44). A plasmid encoding a soluble, catalytically inactive huACE2 ectodomain (residues 1–740) with a C-terminal His tag, cloned into the pCAGGS expression vector, has been previously described (12). Plasmids encoding wild-type huACE2 and 18 naturally occurring huACE2 variants—S19P, K26R, K26E, T27A, E35K, E37K, K68E, M82I, P84T, E329G, D355N, P389H, P426A, D427Y, R559S, S692P, N720D and L731F—were previously cloned into the pCSIB expression vector (12).

### VHH expression plasmids

Monomeric VHHs constructs — For binding characterization, epitope binning experiments, and antiviral assays with monomeric VHHs, DNA sequences encoding the identified VHHs (Table S1) from either discovery pipeline were synthesized gene fragments (Integrated DNA Technologies) and inserted into a bacterial pET21-pelB expression vector bearing an N-terminal pelB leader sequence and C-terminal 6×His tag, via BamHI and XhoI restriction sites and T4 ligation. Cloning, protein expression and purification followed the methods described in (24, 47).

Dimeric VHH constructs— All dimeric VHHs and Fc fusion constructs used in antiviral assays and other experiments in this study were produced using a mammalian Expi293 cell-based expression system (Thermo Fisher Scientific; A40002926). To generate VHH dimers, we used the pCAGGS expression vector—containing a multiple cloning site (MCS) with EcoRI/NotI—as the backbone. After linearization with EcoRI and NotI, the vector was assembled by NEBuilder HiFi DNA Assembly to contain the following tandem elements: EcoRI – murine immunoglobulin signal peptide sequence (GenBank accession no. DQ407610) – AgeI – VHH seq #1 – SalI – (GGSGGGSGGGGSGG) – BamHI – VHH seq #2 – XhoI – (GGGSGG) – His₉* – NotI. Subsequently, individual VHH monomers were cloned into either the N-terminal or C-terminal position by T4 ligation, using the AgeI/SalI or BamHI/XhoI restriction site pairs, respectively.

Bivalent VHH-Fc fusion constructs— To generate bivalent Fc fusion constructs, we excised the region encoding the heavy chain and Fc constant domains from the antibody expression vector described in (12, 50) using AgeI and HindIII. The excised region was substituted with a cassette that, in sequence, contains: AgeI – VHH – BamHI – (GlySer) – GlyGly – human IgG1 Fc (LALA/LS) – HindIII. The Fc domain is derived from human IgG1 and includes the L234A/L235A (LALA) substitutions, which abolish Fcγ receptor interaction, and the M428L/N434S (LS) substitutions, which enhance binding to the neonatal Fc receptor (FcRn), thereby prolonging antibody half-life *in vivo* (12, 51–53). This VHH-Fc vector served as a backbone for subsequent constructs. Individual VHH sequences were PCR-amplified using primers with flanking AgeI and BamHI cut sites and then ligated into the vector with T4 DNA ligase (New England Biolabs).

Tetravalent VHH–Fc fusions— To generate tetravalent Fc constructs, VHH dimers were PCR-amplified from the corresponding pCAGGS plasmids using primers that included flanking regions with the AgeI and BamHI restriction sites present in the Fc backbone vector, and the resulting fragments were assembled by T4 ligation.

### Protein and VHH expression and purification

Nanobody monomers were purified by periplasmic expression in E. coli and His-tag purification as previously described, with endotoxin removed using Triton X-114 (22, 47). All other recombinant proteins were produced using the Expi293 mammalian expression system (Thermo Fisher Scientific; A40003643). Cells were transfected with the appropriate expression plasmids using ExpiFectamine 293 according to the manufacturer’s instructions (Thermo Fisher Scientific; A40002926). Culture supernatants were harvested three to four days post-transfection and clarified by filtration through 0.22 μm membrane filters. Clarified supernatants intended for production of monomeric His-tagged huACE2 soluble ectodomain constructs (used as immunogen or capture antigen for the YD) and dimeric VHHs were loaded onto Ni-NTA agarose (Qiagen), washed extensively, and eluted with 200 mM imidazole in PBS. VHH dimers were similarly processed. Bivalent and tetravalent Fc-fusion constructs were purified from clarified supernatants by overnight incubation with Protein G Sepharose 4 Fast Flow (Cytiva) at 4 °C. The resin was packed into a column and washed; bound protein was eluted with 0.1 M glycine, pH 2.9, into tubes containing 1/10th volume of 1 M Tris, pH 8.0, for immediate neutralization. All purified recombinant proteins, including Fc constructs, were dialyzed against PBS prior to use in downstream experiments.

### Nanobody binding characterization and epitope binning

Nanobody candidates were screened by ELISA to assess binding affinity. Recombinant huACE2 was immobilized on MaxiSorp plates overnight at 4°C at 5 µg/ml in PBS. Plates were washed 3x with PBST (PBS + 0.1% Tween-20) between all subsequent steps. After 1 hr blocking with 10% nonfat milk in PBS, purified nanobodies were diluted in PBST to 0.8, 4, 20, and 100 nM, and incubated for 1hr at room temperature. HRP-conjugated anti-VHH secondary antibody (Jackson ImmunoResearch) was diluted 1:10,000 in PBST + 5% milk and incubated for 1 hr at room temperature. Signal was developed with TMB substrate (Thermo Fisher Scientific) and A450 absorbances measured. EC50s were estimated by fitting a 4-parameter logistic curve.

Binding kinetics (k_a_, k_d_, and K_D_) of selected anti-huACE2 nanobodies were obtained on a Biacore 8K instrument (Cytiva). Recombinant huACE2 was immobilized on a Series S CM5 sensor chip at 5 µg/ml in 10mM sodium acetate, pH 4.5 using EDC/NHS coupling chemistry according to the manufacturer’s guidelines. Nanobodies and RBD were prepared as analytes and run in buffer containing 20 mM HEPES pH 7.4, 150 mM NaCl, and 0.05% Tween-20. Analytes were injected at 30 µl/min in parallel kinetics experiments at concentrations of 0.14, 0.41, 1.23, 3.70, 11.11, 33.33, and 100 nM, with an association time of 180 sec and a dissociation time of 1200 sec. Residual bound protein was removed between experiments using 10 mM glycine-HCl pH 3.0 + 1M MgCl_2_. Data were analyzed using the Biacore Insight Evaluation software, fitting a Langmuir 1:1 binding model to sensorgrams to calculate kinetic parameters.

For epitope binning, pairs of nanobody or RBD proteins were sequentially flowed over immobilized huACE2 using Biacore tandem dual injections according to the manufacturer’s guidelines. Proteins were injected at concentrations of 200 nM with a flow rate of 10 µl/min. Contact time for the first protein was 120 sec, followed by 150 sec for the second protein, then a 30 sec dissociation. Response signal for the second antibody was measured in a 10 sec window at the beginning of dissociation. The chip was regenerated between experiments as above. Data were analyzed using the Biacore software epitope binning module.

### Structural prediction of nanobody complexes

The AlphaRED pipeline was used for sequence-based structural prediction of nanobody:huACE2 complexes, based on AlphaFold predictions followed by replica exchange docking, and performed on the ROSIE online server (26, 54). Initial confidence scoring was based on AlphaFold pLDDT and PAE metrics, as well as Rosetta interface scores and RMSDs for top docking models. ipSAE was also used to assess the AlphaFold prediction (55).

### Sarbecovirus spike-pseudotyped virus production and infectivity inhibition assay

Pseudotyped HIV-1 virions displaying sarbecovirus spike proteins were generated as previously described (12). Briefly, 1 × 10⁷ HEK 293T cells plated in a 15 cm dish were co-transfected with 25 μg of an envelope-deficient HIV-1 proviral plasmid encoding NanoLuc and 7.5 μg of the matching spike expression plasmid. The following morning, cells were washed twice with PBS; culture supernatants were harvested 48 h post-transfection, filtered (0.22 μm), and concentrated using Lenti-X Concentrator (TaKaRa). To titrate pseudovirus infectivity, viral stocks were serially diluted five-fold and added to Huh-7.5 cells, seeded one day earlier in 96-well black plates, that endogenously express huACE2 and are permissive to SARS-CoV-2 entry (12). Forty-eight hours post-infection, cells were lysed and NanoLuc luciferase activity was quantified using the Nano-Glo Luciferase Assay System on a GloMax Navigator Microplate Luminometer (Promega). Luminescence is reported as relative light units (RLU).

To assess antiviral activity, anti-huACE2 VHH constructs were serially diluted in fivefold steps, in triplicate (starting concentration, 3 μg ml⁻¹; seven dilutions) in 96-well plates and incubated with Huh-7.5 target cells for 1 h at 37 °C. The anti-huACE2 monoclonal antibody 5B4 served as the positive control, while the irrelevant anti-HIV-1 monoclonal antibody 3BNC117 served as the negative control (12, 39). Following the pre-incubation period, treated cells were infected with the corresponding spike-pseudotyped viruses at a dose to achieve between 10^6^-10^7^ RLUs, measured at 48 h after infection, as described above. Values were normalized to infected, untreated positive control (set to 100%) and uninfected negative control (set to 0%). Normalized values were fitted by nonlinear regression using the “[inhibitor] vs. normalized response” model in GraphPad Prism, with top and bottom constrained to 100% and 0%, respectively. IC50 values were interpolated as the concentration producing 50% inhibition. All antiviral assays were performed in three independent experiments with three technical replicates each, except the initial screening experiments of VHHs that were performed once in duplicates.

### Flow cytometric analysis of cell-surface huACE2 binding by anti-huACE2 VHH-Fc fusions

To evaluate anti-huACE2 VHH-Fc fusion binding to cell-surface ACE2, A549 human alveolar epithelial cells engineered to stably express either huACE2 or mouACE2 with the C-term 3xHA tag were utilized. A549 cells stably expressing hu or mouACE2 and parental A549 cells were detached with 10 mM EDTA in PBS and 2 × 10⁵ cells were incubated with each VHH-Fc was at at 2.5, 0.5, or 0.1 μg·ml⁻¹ for 2 h at 4 °C. The unrelated anti-HIV-1 mAb 3BNC117 was included as a negative control and the anti-huACE2 mAb 5B4 as a positive control (12, 39). After washing, Fc domains bound to the cell surface were then detected by staining with an Alexa Fluor 488-conjugated goat anti-human IgG secondary antibody (Thermo Fisher Scientific) and analyzed using an Attune NxT Acoustic Focusing Cytometer (Thermo Fisher Scientific). Parental (unmodified) A549 cells were processed in parallel to control for non-specific cell-surface binding. To account for ACE2 expression levels, cells were fixed in parallel, permeabilized and stained with an anti-HA.11 epitope tag mouse mAb conjugated with PE/Dazzle 594 (Biolegened; clone 16B12).

To assess the impact of naturally occurring huACE2 polymorphisms on VHH-Fc binding, HEK 293T cells were transiently transfected with plasmids encoding either wild-type huACE2 or one of 18 huACE2 variants (S19P, K26R, K26E, T27A, E35K, E37K, K68E, M82I, P84T, E329G, D355N, P389H, P426A, D427Y, R559S, S692P, N720D, or L731F); each construct was co-transfected with an mCherry-expressing pCR3.1 plasmid as a transfection control in a 6-well dish format. Cells transfected with only the mCherry plasmid served as the negative control. Two days post-transfection, cells were harvested with EDTA, then incubated with anti-huACE2 VHH-Fc fusions (2 μg ml⁻¹) and stained with Alexa Fluor 488-conjugated goat anti-human IgG mAb (Thermo Fisher Scientific). The mCherry-positive population was gated to quantify anti-huACE2 VHH-Fc binding, which was measured as geometric mean fluorescence intensity.

### huACE2 enzymatic activity assay

Endogenous huACE2 catalytic activity in the presence of anti-huACE2 VHH-Fc fusions was evaluated using the fluorogenic substrate Mca-APK(Dnp) (AnaSpec) (56). Briefly, Mc-Ala fluorescence, normally quenched by Dnp, is restored when huACE2 cleaves the peptide and the two moieties separate; fluorescence is monitored at 330/390 nm and used to quantify enzymatic activity. Huh-7.5 cells were seeded at 1 × 10⁴ per well in black-walled, 96-well plates and maintained for 48 h in 200 μL of complete medium. Cells were washed once and incubated for 1 h at 37 °C with 40 μL of VHH-Fc fusions, mAb 5B4 (positive control), 3BNC117 (negative control) — at the same serial dilution concentrations used in the antiviral assays — or with the ACE2-specific small-molecule inhibitor MLN-4760 (Tocris Bioscience; starting concentration: 20 μM); all reagents were diluted in phenol red-free DMEM (Gibco). Subsequently, 40 μL of Mca-APK(Dnp) substrate diluted in PBS was added, and the reaction proceeded for 30 min at 37 °C. Wells containing substrate only and wells containing medium only served as the blank and positive controls, respectively. Fluorescence from substrate cleavage and Mca-Ala release was measured at excitation/emission wavelengths of 330/390 nm using a CLARIOstar Plus Microplate Reader (BMG Labtech). ACE2 enzymatic activity was quantified by fluorogenic substrate cleavage and relative fluorescence units expressed as a percentage of the uninhibited (no-inhibitor) control, set to 100%.

### huACE2 compartmentalization assay

To determine whether anti-huACE2 VHH-Fc fusions alter huACE2 subcellular localization, live A549 cells stably expressing huACE2 carrying an N-terminal extracellular HA epitope tag (described in (12)) were incubated with the lead anti-huACE2 construct Nb15-7-Fc (1 μg ml⁻¹) or the irrelevant control mAb 3BNC117 for 1h at 4 °C (surface-restricted) or 37 °C (endocytosis-permissive), as previously described (12). Cells were then washed, fixed in 4% paraformaldehyde in PBS, quenched with 10 mM glycine, and permeabilized with 0.1% Triton X-100. Total huACE2-HA was detected with mouse anti-HA.11 antibody (clone 16B12; BioLegend; cat. no. 901503; 1:1000). Bound antibodies were visualized with goat anti-mouse Alexa Fluor 594 (1:500; detects HA-tagged huACE2) and/or goat anti-human Alexa Fluor 488 (1:500; detects the VHH-Fc fusion) (Thermo Fisher Scientific). As a positive control for receptor-mediated endocytosis, cells were stained with a anti-CD44 antibody (clone F10-44-2; Abcam; cat. no. ab30405; dilution 1:20) detected with a secondary goat anti-rabbit Alexa Fluor 488 (1:500). Images were acquired using a DeltaVision OMX SR structured illumination microscope (GE Healthcare) and were processed in Fiji. For each condition, five cells were randomly selected from single optical sections for quantification of intracellular huACE2-HA fluorescence. In Fiji, a line profile traversing the full cell diameter (from plasma membrane, through the nucleus, to the opposite plasma membrane) was drawn for each cell; fluorescence intensity values from Plot Profiles for the Alexa Fluor 488 or Alexa Fluor 594 channel were then extracted. Each profile was normalized to 100% of the cell diameter to account for differences in absolute distances between cells, and the five normalized profiles per condition were subsequently overlaid and plotted.

### Animal experiments

Female C57BL/6J (B6) mice (strain #000664), huACE2 knock-in (KI) mice where huACE2 cDNA replaces the endogenous mouse *Ace2* locus (B6.129S2(Cg)-Ace2^tm1(ACE2)Dwnt/J, strain 035000) and HuACE2-K18 transgenic mice (B6.Cg-Tg(K18-ACE2)2Prlmn/J, strain 034860) that express huACE2 under the cytokeratin-18 promoter were obtained from the Jackson Laboratory (40). huACE2-K18 mice were crossed with IFNR^−/−^ mice to generate huACE2-K18-IFNR^−/−^ transgenic mice (41, 57). Mice were 10-12 weeks old at the start of experiments and were housed at 22 °C with 30– 70% humidity on a 12 h light/12 h dark cycle, with *ad libitum* access to food and water; they were acclimatized for at least 2 weeks before experimental procedures. All animal procedures were conducted in accordance with protocols approved by the Rockefeller University Institutional Animal Care and Use Committee. Experiments involving replication-competent SARS-CoV-2 were performed under animal biosafety level biosafety level 3 (ABSL-3) conditions.

### Analysis of anti-huACE2 VHH-Fc fusion pharmacokinetics in mice

B6 and huACE2 KI mice (n = 3 per group) received a single subcutaneous injection on day 0 of 1.7 nmol per mouse of either NbA15-7-Fc or NbA15-117-Fc to determine the pharmacokinetic properties of the anti-huACE2 VHH-Fc fusions; this dose is the molar equivalent of 250 μg of a conventional IgG1 (approximately 12.5 mg kg⁻¹). On days 1, 4, and 7 after injection, blood samples were collected into Microvette CB 300 Serum tubes (Sarstedt). We quantified serum concentrations of the injected VHH-Fc constructs by means of the functional pseudovirus neutralization assay described above (see “Sarbecovirus spike-pseudotyped virus production and infectivity inhibition assay”). Mouse sera were serially diluted (starting at 4 µL), pre-incubated with Huh-7.5 target cells for 1 h at 37 °C, and then infected with SARS-CoV-2 Wuhan-Hu-1 spike-pseudotyped HIV-1 NanoLuc reporter virions; NanoLuc luciferase activity was measured 48 h post-infection. Values were normalized to infected, untreated positive control (set to 100%) and uninfected negative control (set to 0%). Known concentrations of each VHH-Fc fusion protein were serially diluted with equivalent volumes of control mouse serum and assayed in parallel with the mouse serum samples to construct standard calibration curves, permitting interpolation of serum VHH-Fc concentrations from the dose–response curves. Serum half-lives were calculated by fitting the concentration–time data to a one-phase exponential decay model using GraphPad Prism (Version 11.0.0). During the pharmacokinetic study and all subsequent in vivo experiments, mice were monitored daily for clinical signs of adverse effects, including changes in body weight, activity, grooming, and posture.

### Prophylactic efficacy against SARS-CoV-2 in huACE2 knock-in mice

To evaluate prophylactic efficacy, huACE2 KI mice (n = 10 per group) received a single intraperitoneal injection of 1.7 nmol NbA15-7-Fc or PBS (vehicle control) two days before intranasal challenge with 2 × 10⁵ plaque-forming units (PFU) of SARS-CoV-2, strain USA-WA1/2020, administered in 30 μL under deep anesthesia with inhalant Isoflurane administered at 3-4% in oxygen. At 3 days post-infection, the lungs were harvested where each lung lobe (right and left) were homogenized and processed separately and RNA was extracted using TRIzol and phase separation using chloroform. Genomic RNA was quantified by RT-qPCR targeting the N gene using 1-Step Kit, PowerSYBR Green RNA-to-CT (ThermoFisher #4389986) and StepOne Plus Real-Time PCR system (Applied Biosystems), where each lung lobe was considered as replicate (12). The primers used were Integrated DNA Technologies 2019-nCOV_N1 Forward Primer: 5′-GACCCCAAAATCAGCGAAAT-3′ Aliquot, 50nmol, catalog #10006821, and Integrated DNA Technologies 2019-nCOV_N1 Reverese Primer: 5′-TCTGGTTACTGCCAGTTGAATCTG-3′ Aliquot, 50nmol, catalog #10006822. Viral RNA copies were normalized to micrograms of total RNA with use of Integrated DNA Technologies (IDT) 2019-nCOV_N_Positive Control, catalog #10006625. The limit of detection was 100 copies per μg RNA. To confirm proper i.p. administration of our modalities, serum was collected from the submandibular vein before virus challenge and tested for comparable antiviral activity. An RT-qPCR reaction for a housekeeping gene (GAPDH; IDT) was run in parallel to ensure comparable RNA concentrations among reactions.

### Assessment of prophylactic efficacy against VSV/SARS**-**CoV in K18-huACE2 IFNAR⁻/⁻ mice

To assess the breadth of protection conferred by anti-huACE2 VHH-Fc fusions against related sarbecoviruses, K18-huACE2 IFNAR⁻/⁻ transgenic mice (n = 6 per group) received intraperitoneal injections of 1.7 nmol NbA15-7-Fc or PBS two days prior to intranasal challenge with 2×10^4^ PFU of a replication-competent VSV pseudotyped with the SARS-CoV spike protein as described in (44). Viral load was quantified as described above. The primers used were 5′-CTCTGCCGACTTGGCACAAC-3’ and 5′-TTCAAACCATCCGAGCCATTCG-3’, targeting RNA sequences that encode the nucleoprotein (N) of VSV. A reference plasmid, containing VSV N, was used as a standard to interpolate the number of viral RNA copies per microgram of input lung RNA in each reaction.

### Animal ethics statement

Llama care was performed at the USDA regulated Capralogics, Inc. research facility 14-R-0156 and under OLAW Assurance number D16-00582 (A4079-01) with all protocols, standard operation procedures and animal care procedures at this facility performed in accordance with the Public Health Service (PHS) Policy on the Humane Care and Use of Laboratory Animals, the Animal Welfare Act and the 8^th^ Edition of The Guide for the Care and Use of Laboratory Animals as well as the AVMA Guidelines for the Euthanasia of Animals: 2020 Edition.

Mouse experiments were performed at the animal facility at Rockefeller University Collaborative Bioscience Center. All procedures have been approved under IACUC protocol 21016-H, approved 4/26/24. The center is fully accredited by the Association for Assessment and Accreditation of Laboratory Animal Care International. At no point in these studies were the animals allowed to suffer. The health of all mice was monitored frequently and euthanasia performed following the the American Veterinary Medical Association (AVMA) Guidelines.

## Statistical analysis

Figures were generated using GraphPad Prism (Version 11.0.0). Statistical analysis was conducted using GraphPad Prism (Version 11.0.0).

## Supporting information

ACE2 VHH Supplement

## Acknowledgments

This work was supported by a grant from the Stavros Niarchos Foundation Institute for Global Infectious Disease Research (TH, BTC, MPR, FRC and PDB), by the Howard Hughes Medical Institute (PDB) and the National Institute of Allergy and Infectious Diseases R01 AI189657 (MPR).

## Author Contributions

Conceptualization: ADB, PCF, BTC, MPR, FRC, PDB, TH; Data curation: ADB, RP, PCF, JJ, MA, LS, KRM; Formal analysis: ADB, RP, PCF, JJ, MA; Funding acquisition: BTC, MPR, FRC, PDB, TH; Investigation: ADB, RP, PCF, JJ, MA, VAB, LS, KRM; Methodology: ADB, RP, PCF, JJ, MA, LS, MPR, BTC, FRC, PDB, TH; Project administration: ADB, PCF, BTC, MPR, FRC, PDB, TH; Resources: BTC, MPR, FRC, PDB, TH; Software: ADB, RP, PCF, JJ, MA, LS, KRM; Supervision: ADB, MPR, BTC, FRC, PDB, TH; Validation: ADB, RP, PCF, JJ, MA, VAB, LS, KRM, MPR, BTC, FRC, PDB, TH; Visualization: ADB, RP, PCF, MA, TH; Writing – original draft: ADB, RP, PCF, TH; Writing, reviewing & editing: ADB, RP, PCF, JJ, MA, VAB, LS, KRM, MPR, BTC, FRC, PDB, TH.

