## Supplementary material for "Multivalent Anti-ACE2 Nanobodies Confer Broad Pan-Sarbecovirus Protection": ACE2 VHH Supplement

**Affiliations:** 1) Laboratory of Retrovirology, The Rockefeller University, New York, NY 10065. 2) Laboratory of Cellular and Structural Biology, Rockefeller University, 1230 York Ave, Box 213, New York, NY 10021, USA. 3) Laboratory of Mass Spectrometry and Gaseous Ion Chemistry, The Rockefeller University, New York, USA. 4) Laboratory of Cell Cycle Genetics, The Rockefeller University, New York, NY 10021, USA.

**Running title:** Anti-ACE2 Nanobodies as pan-sarbecovirus therapeutics

Fig S1

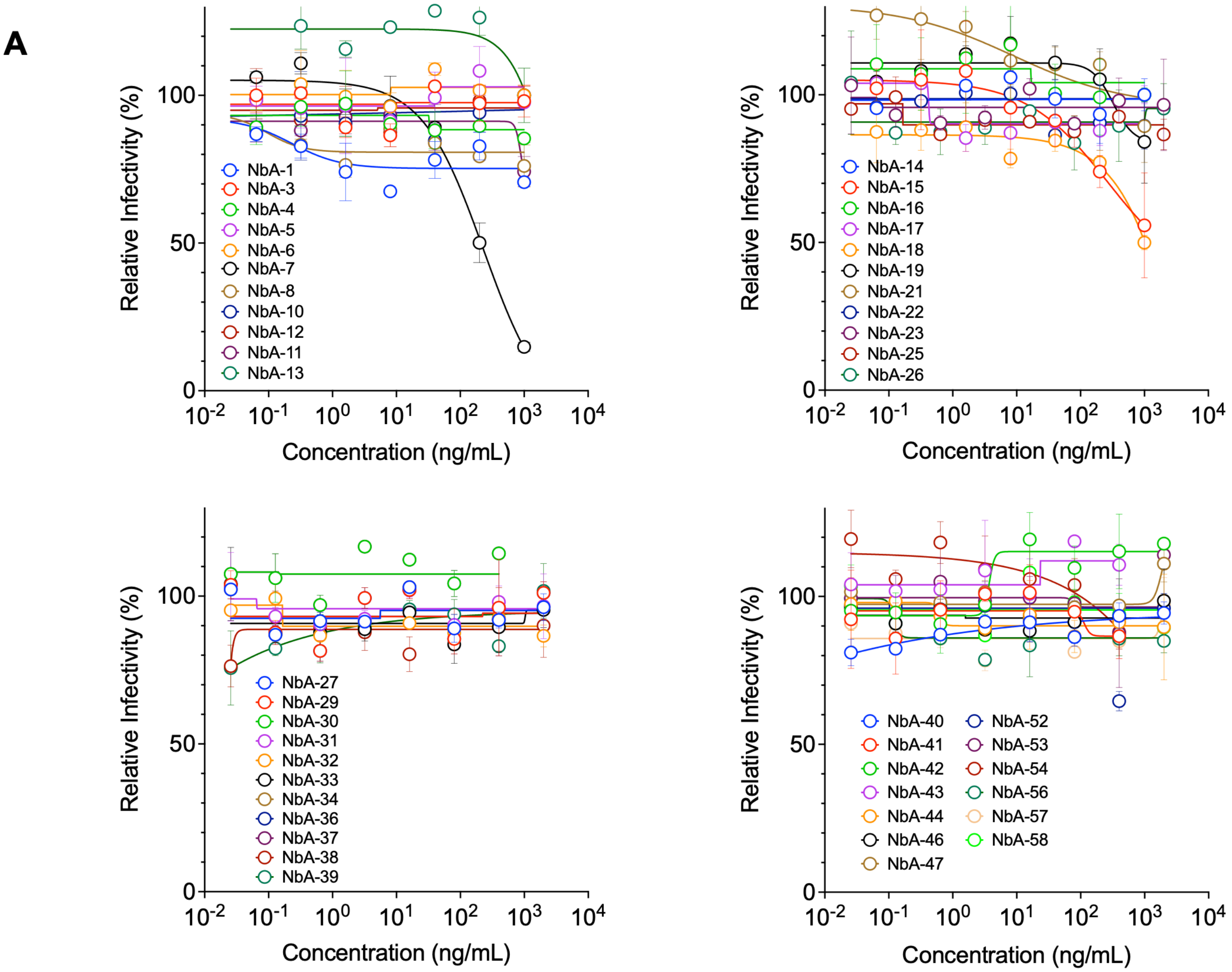
A
Relative Infectivity (%)
Concentration (ng/mL)
NbA-1
NbA-3
NbA-4
NbA-5
NbA-6
NbA-7
NbA-8
NbA-10
NbA-12
NbA-11
NbA-13
NbA-14
NbA-15
NbA-16
NbA-17
NbA-18
NbA-19
NbA-21
NbA-22
NbA-23
NbA-25
NbA-26
NbA-27
NbA-29
NbA-30
NbA-31
NbA-32
NbA-33
NbA-34
NbA-36
NbA-37
NbA-38
NbA-39
NbA-40
NbA-41
NbA-42
NbA-43
NbA-44
NbA-46
NbA-47
NbA-52
NbA-53
NbA-54
NbA-56
NbA-57
NbA-58

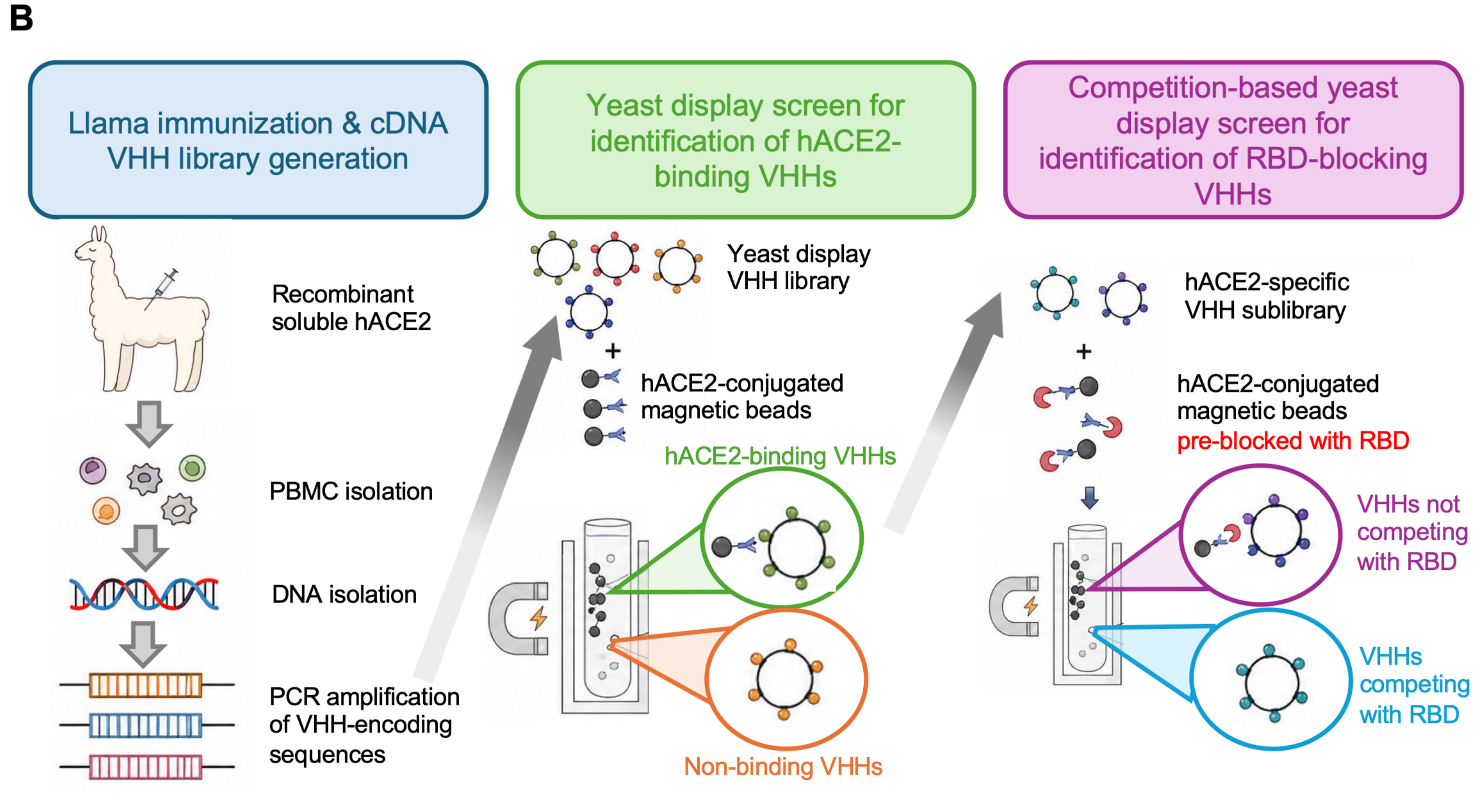
B
Llama immunization & cDNA VHH library generation
Yeast display screen for identification of hACE2-binding VHHs
Competition-based yeast display screen for identification of RBD-blocking VHHs
Recombinant soluble hACE2
PBMC isolation
DNA isolation
PCR amplification of VHH-encoding sequences
Yeast display VHH library
+
hACE2-conjugated magnetic beads
hACE2-binding VHHs
Non-binding VHHs
hACE2-specific VHH sublibrary
+
hACE2-conjugated magnetic beads
pre-blocked with RBD
VHHs not competing with RBD
VHHs competing with RBD

**Figure S1**: **Antiviral activity of anti-huACE2 VHH identified with MS and workflow of a competition-based yeast-display VHH screening method.**

**(A)** Antiviral activity of anti-huACE2 VHHs. Anti-huACE2 VHHs, confirmed to bind ACE2 via ELISA, were serially diluted and incubated with Huh7.5 target cells. Cells were subsequently infected with a SARS-CoV-2-pseudotyped NanoLuc luciferase HIV-1 reporter virus and luciferase activity measured 48hrs post-infection. Values were normalized to infected, untreated positive control (set to 100%) and uninfected negative control (set to 0%). Data represent geometric medians and standard deviation from two independent experiments. **(B)** Schematic workflow for the identification of huACE2-binding VHHs with epitope specificity using a yeast display screen. The screening approach included 3 phases: (1) Llama immunization with recombinant soluble huACE2 and subsequent VHH library construction via BMMC isolation, cDNA synthesis, and PCR amplification of VHH-encoding sequences; (2) Primary yeast display screening using huACE2-conjugated magnetic beads to enrich for ACE2-binding clones; (3) Secondary competitive screening with huACE2-conjugated magnetic beads pre-bound to SARS-CoV-2 RBD to isolate VHHs that compete with RBD for huACE2 binding.

**Figure S2.**

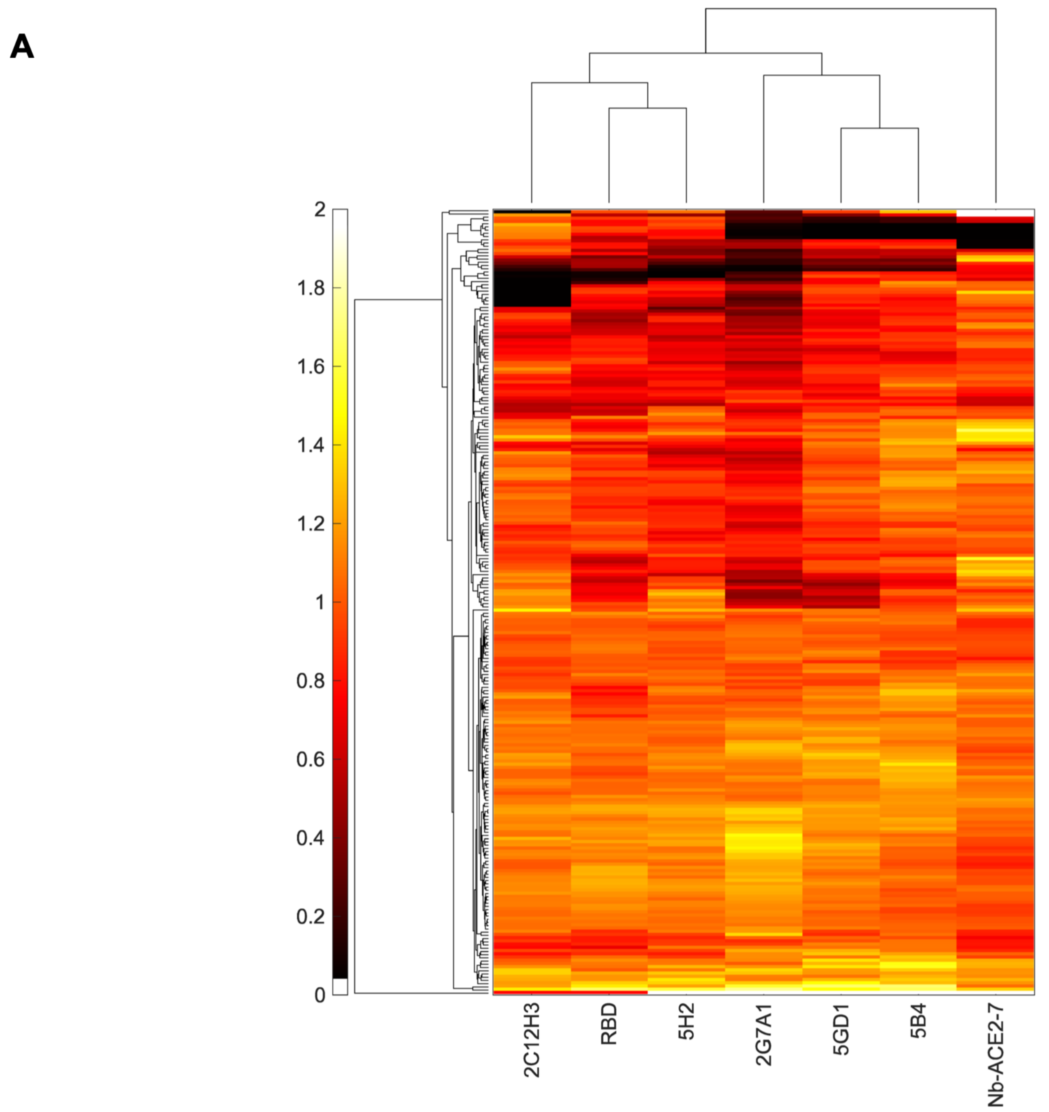
A
2
1.8
1.6
1.4
1.2
1
0.8
0.6
0.4
0.2
0
2C12H3
RBD
5H2
2G7A1
5GD1
5B4
Nb-ACE2-7

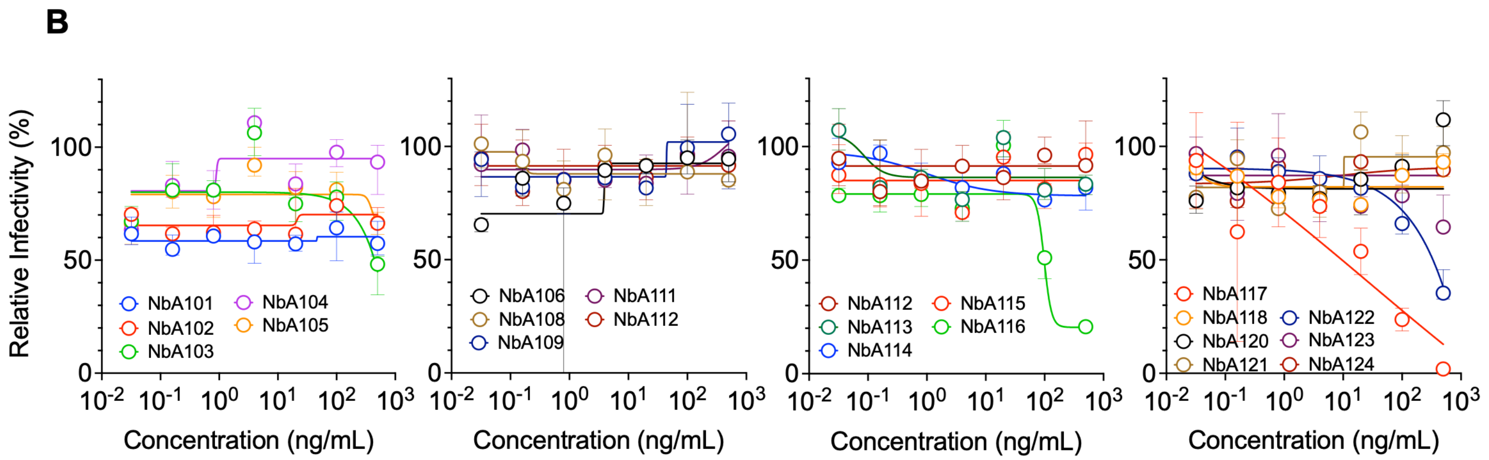
B
Relative Infectivity (%)
NbA101
NbA102
NbA103
NbA104
NbA105
NbA106
NbA108
NbA109
NbA111
NbA112
NbA112
NbA113
NbA114
NbA115
NbA116
NbA117
NbA118
NbA120
NbA121
NbA122
NbA123
NbA124
Concentration (ng/mL)

**Figure S2: Competition-based yeast display screen for identification of anti-huACE2 VHHs and their antiviral activity. (A)** Yeast display competition experiments. Heatmap with hierarchical clustering (MATLAB clustergram) depicting the binding (readcounts) of yeast cells expressing individual VHHs (rows) to huACE2-coated beads pre-saturated with the indicated known binders (columns: SARS-CoV-2 RBD and anti-ACE2 mAbs: 2C1/2H3, 5H2, 2G7A1, 5GD1, 5B4 or NbA-7) normalized to binding to ACE2 in the absence of competitor. For each condition, ACE2-bound beads were first incubated with excess of each known binder, followed by addition of the enriched nanobody yeast display library. Color scale: 0 (black) indicates complete block of binding by competitor, indicating an overlapping epitope; >=1 (yellow-white) indicates no competition and distinct epitopes. Hierarchical clustering was performed on both rows (left dendrogram) and columns (top dendrogram) to group VHH families with similar competition profiles, thereby revealing distinct epitope bins. VHH sequences clustering with the RBD column identify nanobodies targeting the ACE2–RBD interface, representing candidate therapeutic leads for viral neutralization. **(B)** Anti-huACE2 VHHs identified with the yeast display screen were serially diluted and incubated with Huh7.5 target cells that were subsequently infected with a SARS-CoV-2-pseudotyped NanoLuc luciferase HIV-1 reporter virus. Luciferase activity was measured 48hrs post-infection. Values were normalized to infected, untreated positive control (set to 100%) and uninfected negative control (set to 0%). Data represent median and standard deviation from two independent experiments.

**Figure S3.**

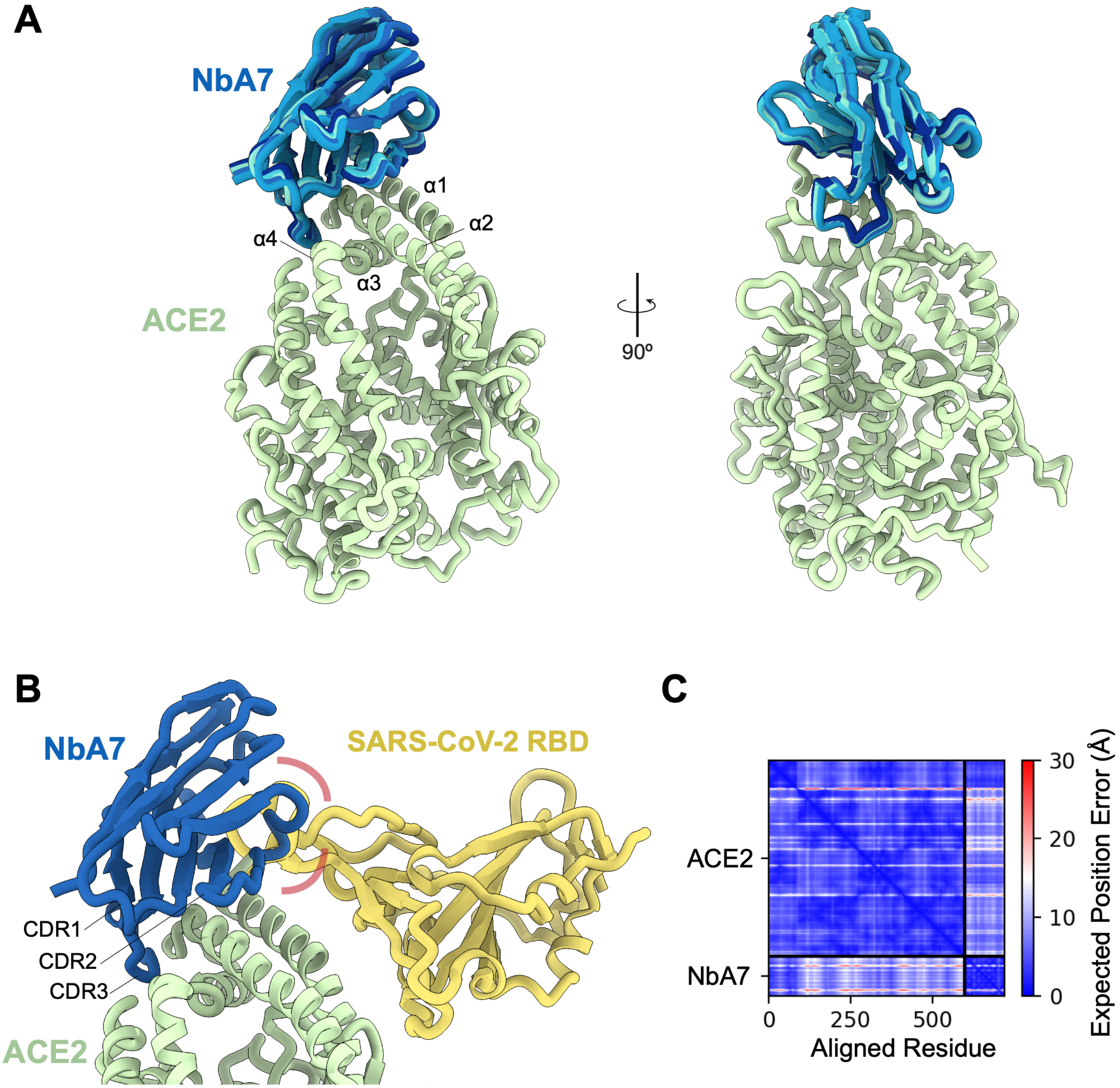
A
NbA7
α1
α2
α4
α3
ACE2
90º
B
NbA7
SARS-CoV-2 RBD
CDR1
CDR2
CDR3
ACE2
C
ACE2
NbA7
0
250
500
Aligned Residue
30
20
10
0
Expected Position Error (Å)

**Figure S3: Structural Prediction of NbA7-ACE2 Interaction and RBD Competition.** **(A)** AlphaFold2-based structural prediction of NbA7 nanobody (blue) in complex with human ACE2 (green), shown in two orientations (90° rotation). ACE2 α-helical domains (α1-α4) are indicated. **(B)** Ternary complex model showing NbA7 (blue), ACE2 (green), and SARS-CoV-2 RBD (yellow) with predicted steric clash region highlighted (pink semicircle). CDR1, CDR2, and CDR3 regions of NbA7 are indicated. **(C)** Predicted aligned error (PAE) matrix for the AlphaFold-Multimer model of the NbA7–ACE2 complex. Axes indicate residue position along the concatenated chains; color indicates the expected positional error (Å) of each residue pair, with off-diagonal blocks reporting confidence in the relative orientation of the two chains.

---

**Figure S4.**

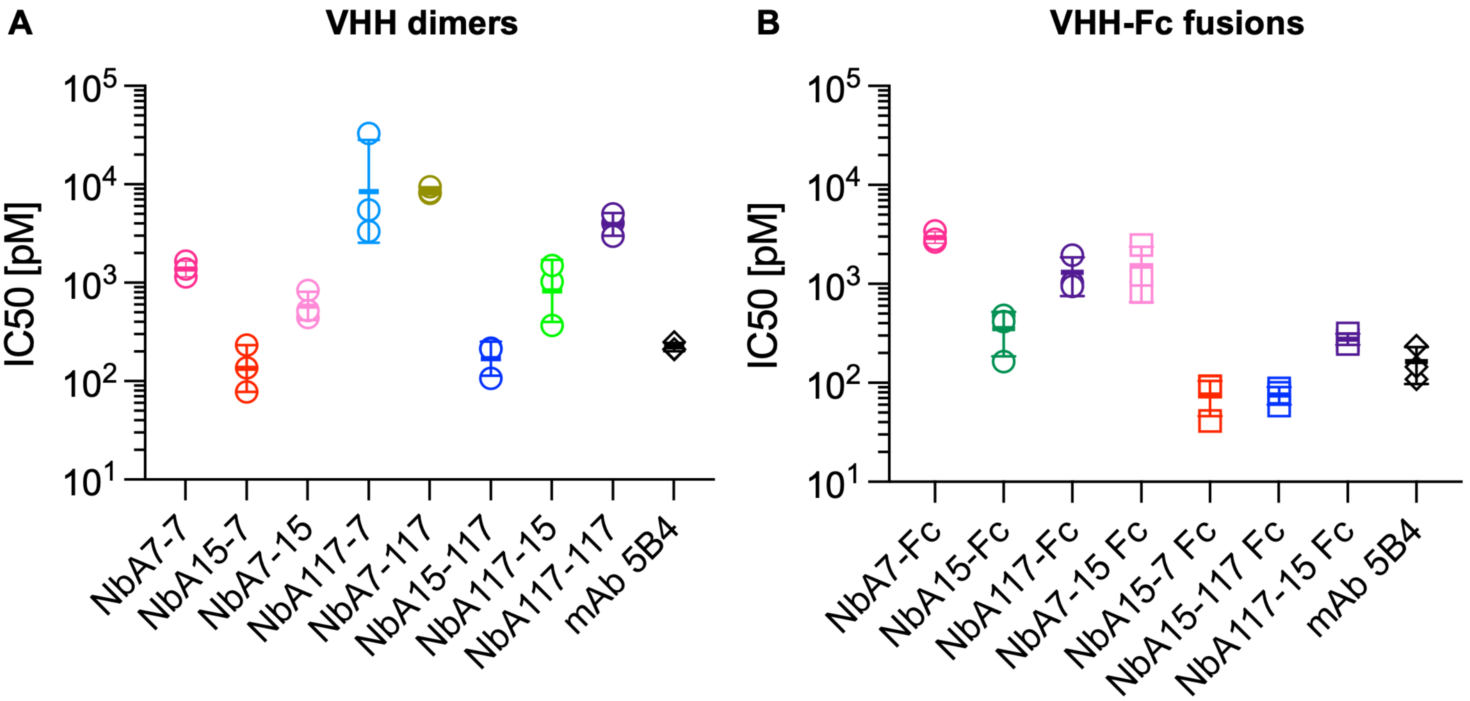

A
VHH dimers
IC50 [pM]
$10^5$
$10^4$
$10^3$
$10^2$
$10^1$
NbA7-7
NbA15-7
NbA7-15
NbA117-7
NbA7-117
NbA15-117
NbA117-15
NbA117-117
mAb 5B4
B
VHH-Fc fusions
IC50 [pM]
$10^5$
$10^4$
$10^3$
$10^2$
$10^1$
NbA7-Fc
NbA15-Fc
NbA117-Fc
NbA7-15 Fc
NbA15-7 Fc
NbA15-117 Fc
NbA117-15 Fc
mAb 5B4

**Figure S4: Dimerization of anti-huACE2 VHHs enhances their antiviral potency.**

**(A)** Dot plot depicting the geometric mean and standard deviation of IC50 values for VHH dimers against SARS-CoV-2 (means of three independent experiments). **(B)** Dot plot depicting the geometric mean and standard deviation of IC50 values for VHH-Fc fusions against SARS-CoV-2 (means of three independent experiments).

---

**Fig S5.**

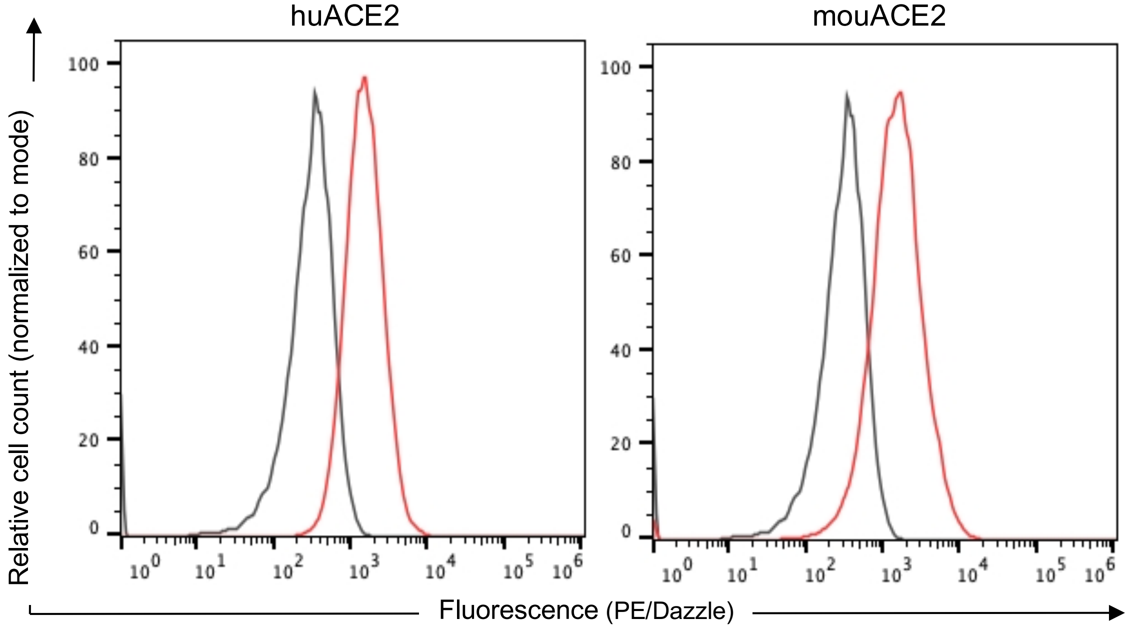


**Fig S5: Expression levels of human and mouse ACE2 evaluated with flow cytometry.** ACE2 expression in A549 cells stably expressing C-terminally HA-tagged huACE2 or mouACE2 (red) compared to parental A549 cells (grey). A549 cells were fixed, permeabilized, incubated in the presence of anti-HA mAb conjugated with PE/Dazzle 594 and analyzed by flow cytometry. Data from a representative experiment are shown.

**Figure S6.**

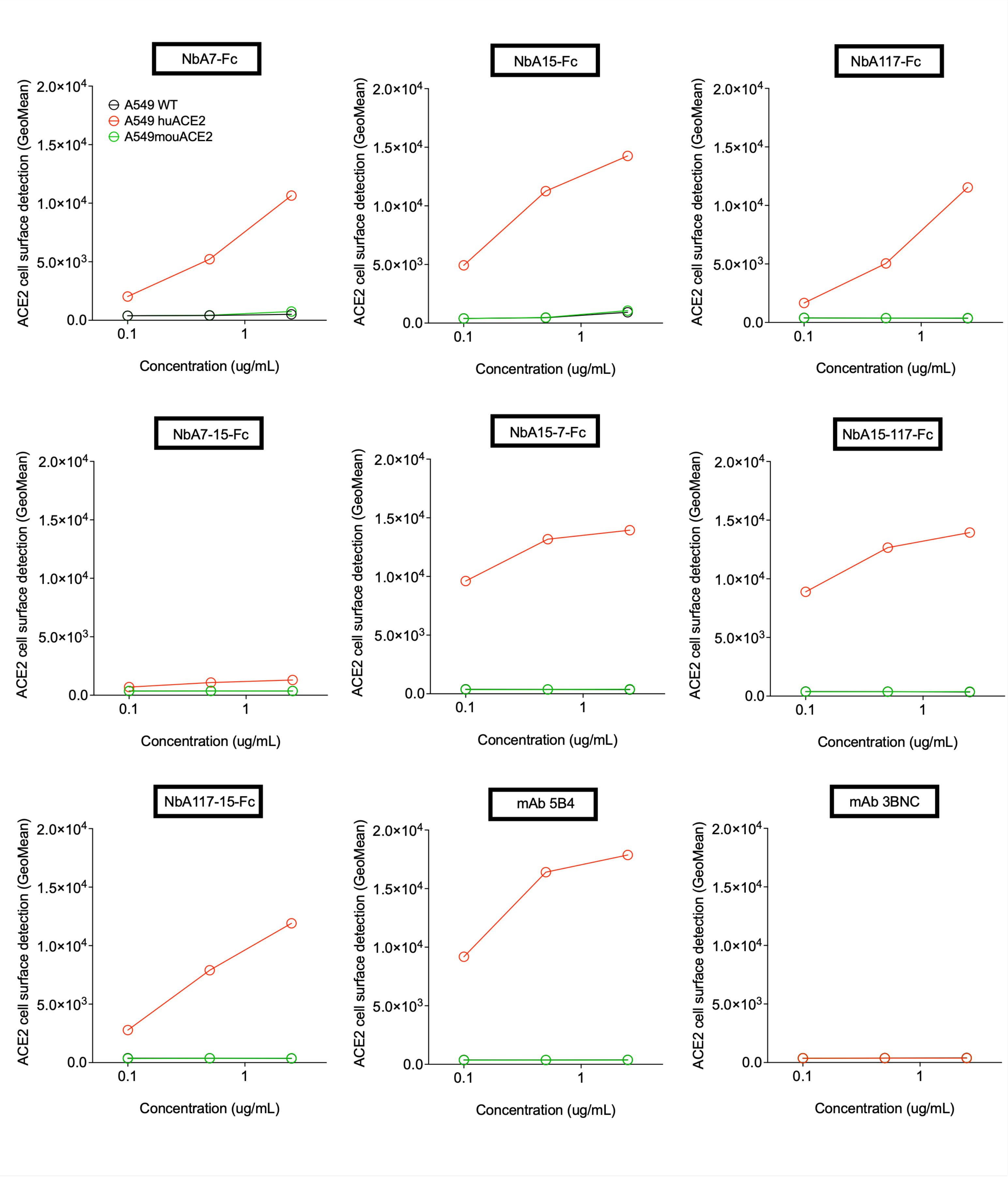
NbA7-Fc
NbA15-Fc
NbA117-Fc
NbA7-15-Fc
NbA15-7-Fc
NbA15-117-Fc
NbA117-15-Fc
mAb 5B4
mAb 3BNC
A549 WT
A549 huACE2
A549mouACE2
ACE2 cell surface detection (GeoMean)
Concentration (ug/mL)
2.0×10^4
1.5×10^4
1.0×10^4
5.0×10^3
0.0
0.1
1

**Figure S6: Cell surface binding of VHH-Fc fusions to ACE2 evaluated with flow cytometry.** Parental A549 cells (blue) or A549 cells stably expressing huACE2 (red) or mouACE2 (green) were incubated in the presence of the indicated anti-hACE2 VHH-Fc fusions, mAb 5B4 (positive control) or irrelevant anti-HIV-1 mAb 3BNC (negative control) at 2.5, 0.5 and 0.1 µg/ml. The cells were then incubated with Alexa Fluor 488 conjugated goat anti-human IgG and analyzed by flow cytometry. Graphs depict the geometric mean fluorescence intensity across different concentrations. Data from a representative experiment are shown.

**Figure S7.**

**A**

**B**

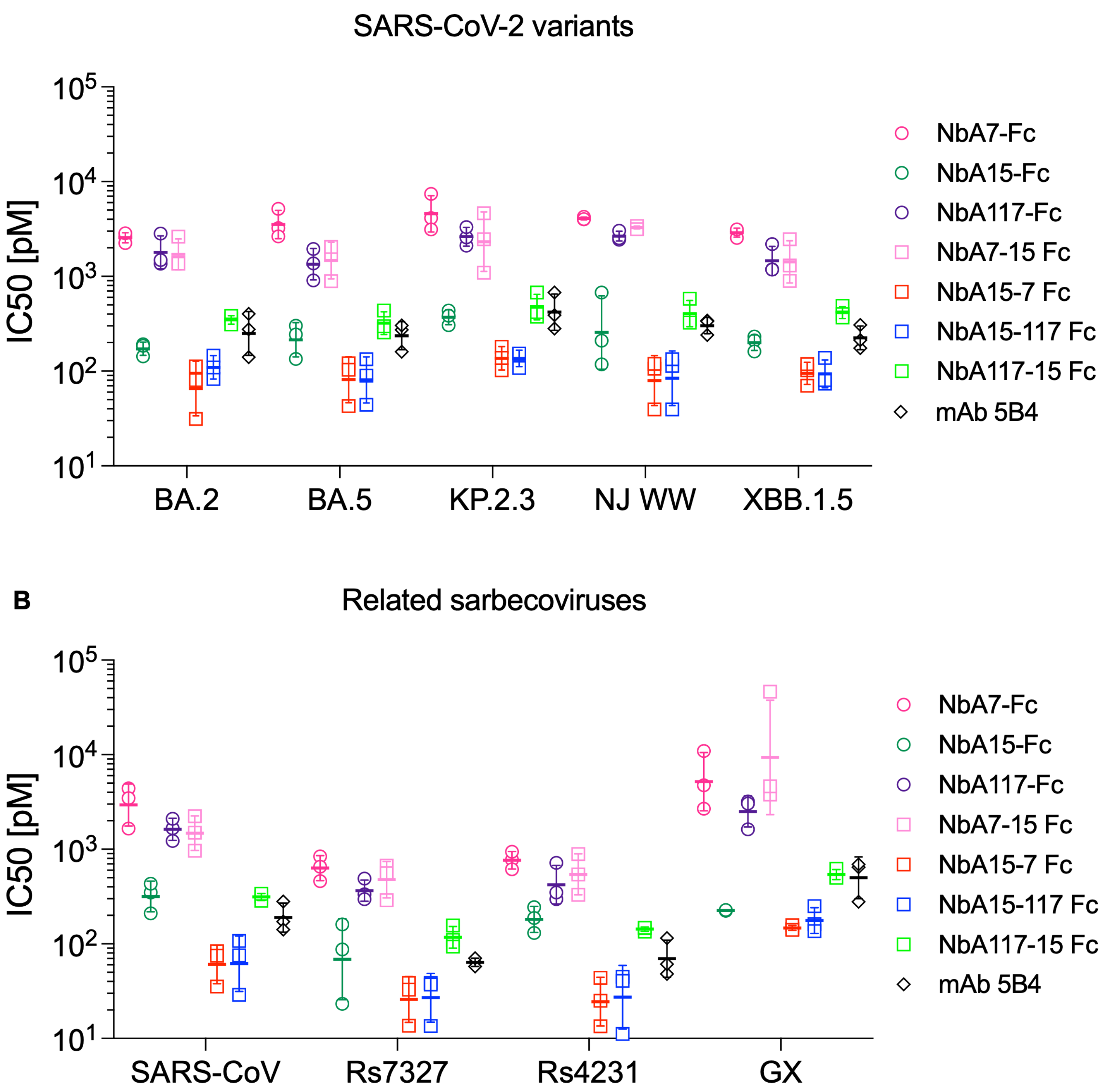
SARS-CoV-2 variants
IC50 [pM]
$10^5$
$10^4$
$10^3$
$10^2$
$10^1$
BA.2
BA.5
KP.2.3
NJ WW
XBB.1.5
NbA7-Fc
NbA15-Fc
NbA117-Fc
NbA7-15 Fc
NbA15-7 Fc
NbA15-117 Fc
NbA117-15 Fc
mAb 5B4
Related sarbecoviruses
IC50 [pM]
$10^5$
$10^4$
$10^3$
$10^2$
$10^1$
SARS-CoV
Rs7327
Rs4231
GX
NbA7-Fc
NbA15-Fc
NbA117-Fc
NbA7-15 Fc
NbA15-7 Fc
NbA15-117 Fc
NbA117-15 Fc
mAb 5B4

**Figure S7: Dimerization of anti-huACE2 VHHs enhances their antiviral potency.**

**(A)** Dot plot depicting the geometric mean and standard deviation of IC50 values for VHH-Fc fusions against SARS-CoV-2 variants (means of three independent experiments). **(B)** Dot plot depicting the geometric mean and standard deviation of IC50 values for VHH-Fc fusions against SARS-CoV and related sarbecoviruses (means of three independent experiments).

Fig S8.

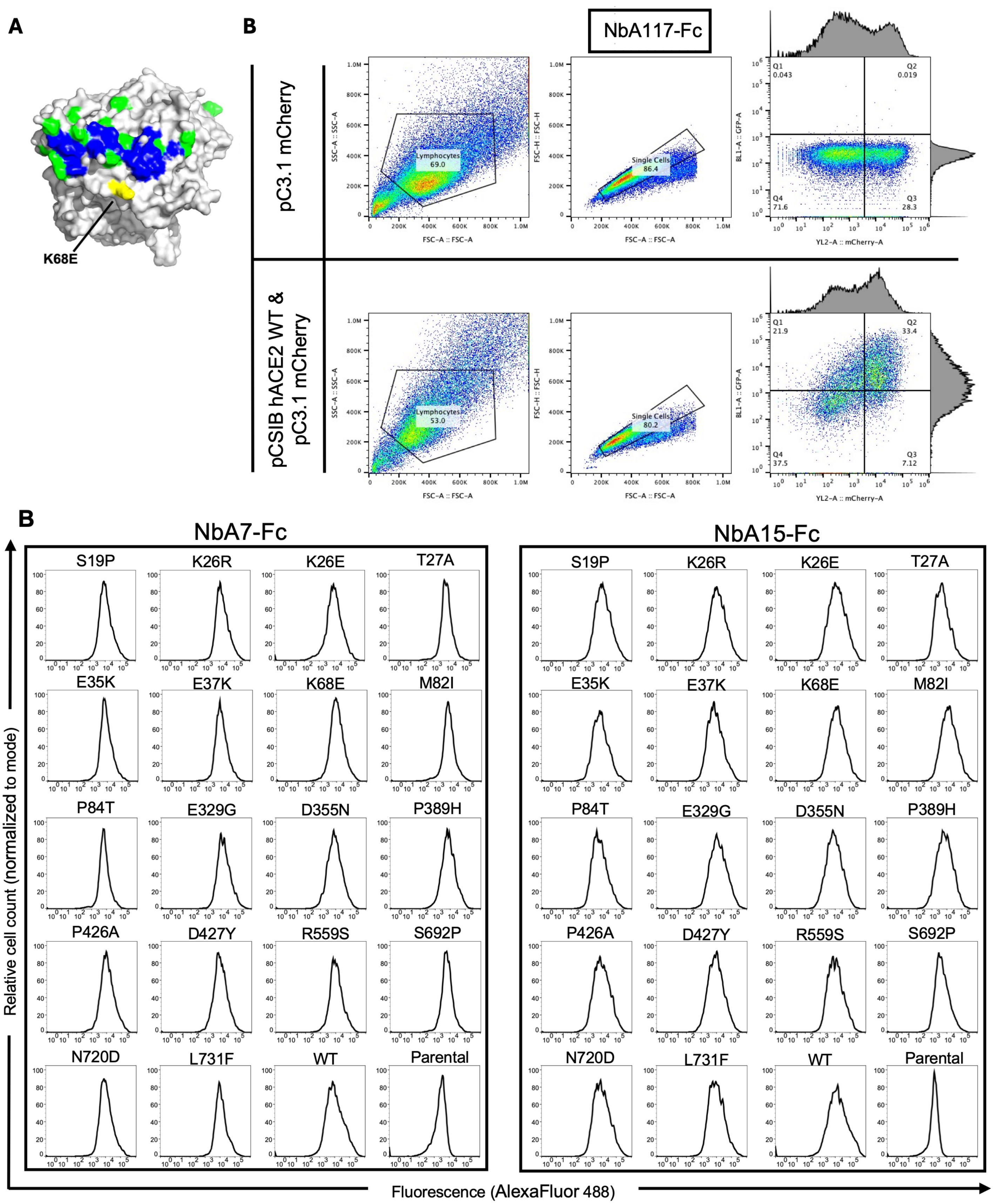
A
K68E
B
NbA117-Fc
pC3.1 mCherry
pCSIB hACE2 WT &
pC3.1 mCherry
Lymphocytes 69.0
Single Cells 86.4
Q1 0.043
Q2 0.019
Q3 28.3
Q4 71.6
Lymphocytes 53.0
Single Cells 80.2
Q1 21.9
Q2 33.4
Q3 7.12
Q4 37.5
SSC-A :: SSC-A
FSC-A :: FSC-A
FSC-H :: FSC-H
BL1-A :: GFP-A
YL2-A :: mCherry-A
B
NbA7-Fc
NbA15-Fc
S19P
K26R
K26E
T27A
E35K
E37K
K68E
M82I
P84T
E329G
D355N
P389H
P426A
D427Y
R559S
S692P
N720D
L731F
WT
Parental
Relative cell count (normalized to mode)
Fluorescence (AlexaFluor 488)

**Fig S8: VHH-Fc Cell Surface Binding to naturally occurring huACE2 mutants.** **(A)** Structure of hACE2 (PDB 6LZG). The SARS-CoV-2 spike binding paratope is colored blue and spatially close amino acids that vary in the human population marked in green. K68 is highlighted in yellow. **(B)** Gating strategy of flow cytometry data comparing NbA117-Fc binding profiles between cells expressing wild type huACE2 or control protein (mCherry). **(C)** Binding of the VHH-Fc fusion protein indicated on cells expressing each of 18 huACE2 mutants. Each histogram shows fluorescence intensity distribution (AlexaFluor 488, x-axis) and relative cell count (y-axis) for individual huACE2 mutants.

**Fig S9.**

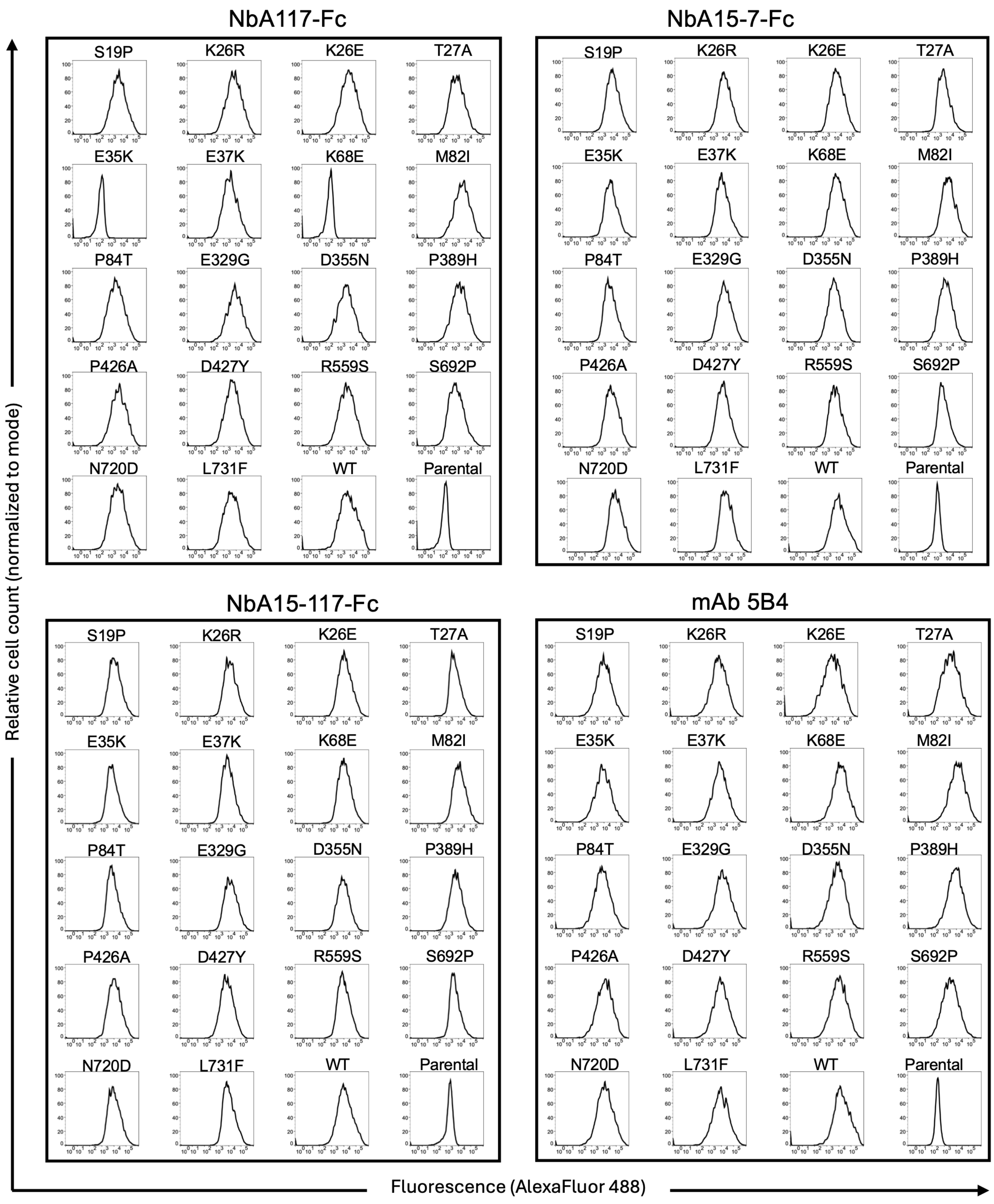

NbA117-Fc
NbA15-7-Fc
NbA15-117-Fc
mAb 5B4
S19P
K26R
K26E
T27A
E35K
E37K
K68E
M82I
P84T
E329G
D355N
P389H
P426A
D427Y
R559S
S692P
N720D
L731F
WT
Parental
Relative cell count (normalized to mode)
Fluorescence (AlexaFluor 488)

**Fig S9: VHH-Fc Cell Surface Binding to naturally occurring huACE2 mutants.** Binding of the VHH-Fc fusion proteins indicated or control anti-huACE2 mAb 5B4 on cells expressing each of 18 huACE2 mutants. Each histogram shows fluorescence intensity distribution (AlexaFluor 488, x-axis) and relative cell count (y-axis) for individual huACE2 mutants.

Fig S10

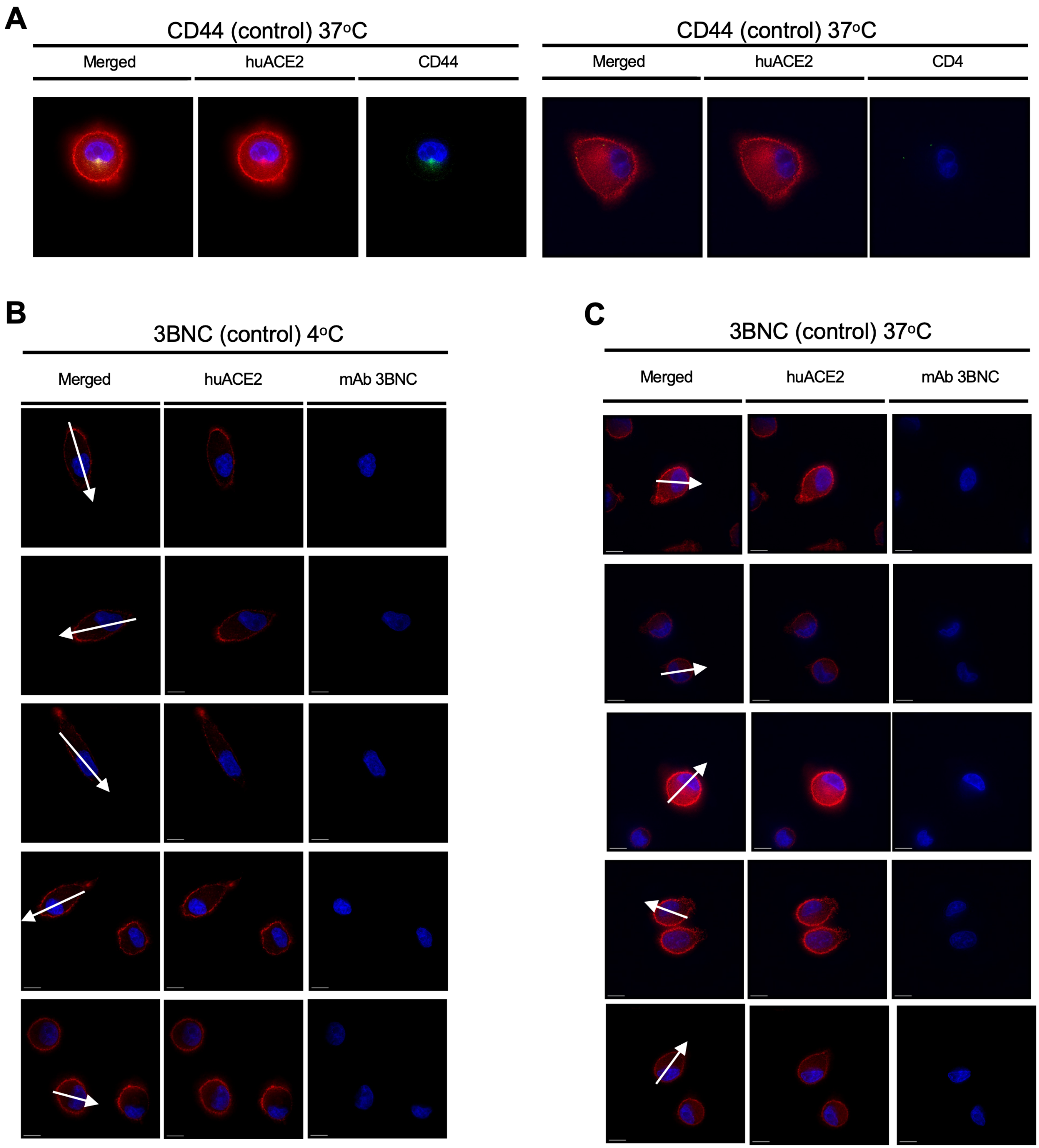
A
CD44 (control) 37oC
Merged
huACE2
CD44
CD44 (control) 37oC
Merged
huACE2
CD4
B
3BNC (control) 4oC
Merged
huACE2
mAb 3BNC
C
3BNC (control) 37oC
Merged
huACE2
mAb 3BNC

**Fig S10: Effect of VHH-Fc fusions on huACE2 localization. (A)** Localization of anti-CD44 antibody (Abcam; ab243894) following incubation with live A549/hACE2-HA cells. Blue stain (DAPI) indicates cell nuclei. Red indicates HA staining of huACE2 (Alexa Fluor™ 594). White arrows represent line profiles traversing the cell diameter for quantification and distribution of fluorescence across each cell. Scale bar: 10 µm. Representative of two independent experiments. **(B-C)** Localization of anti-HIV-1 mAb 3BNC (3BNC117) (Alexa-488, green, right) or huACE2-HA (Alexa-594, red, center) or merged (left) following incubation of live A549/hACE2-HA cells at 4$^{o}$C or 37$^{o}$C as indicated. Additional details as in (A).

**Fig S11**

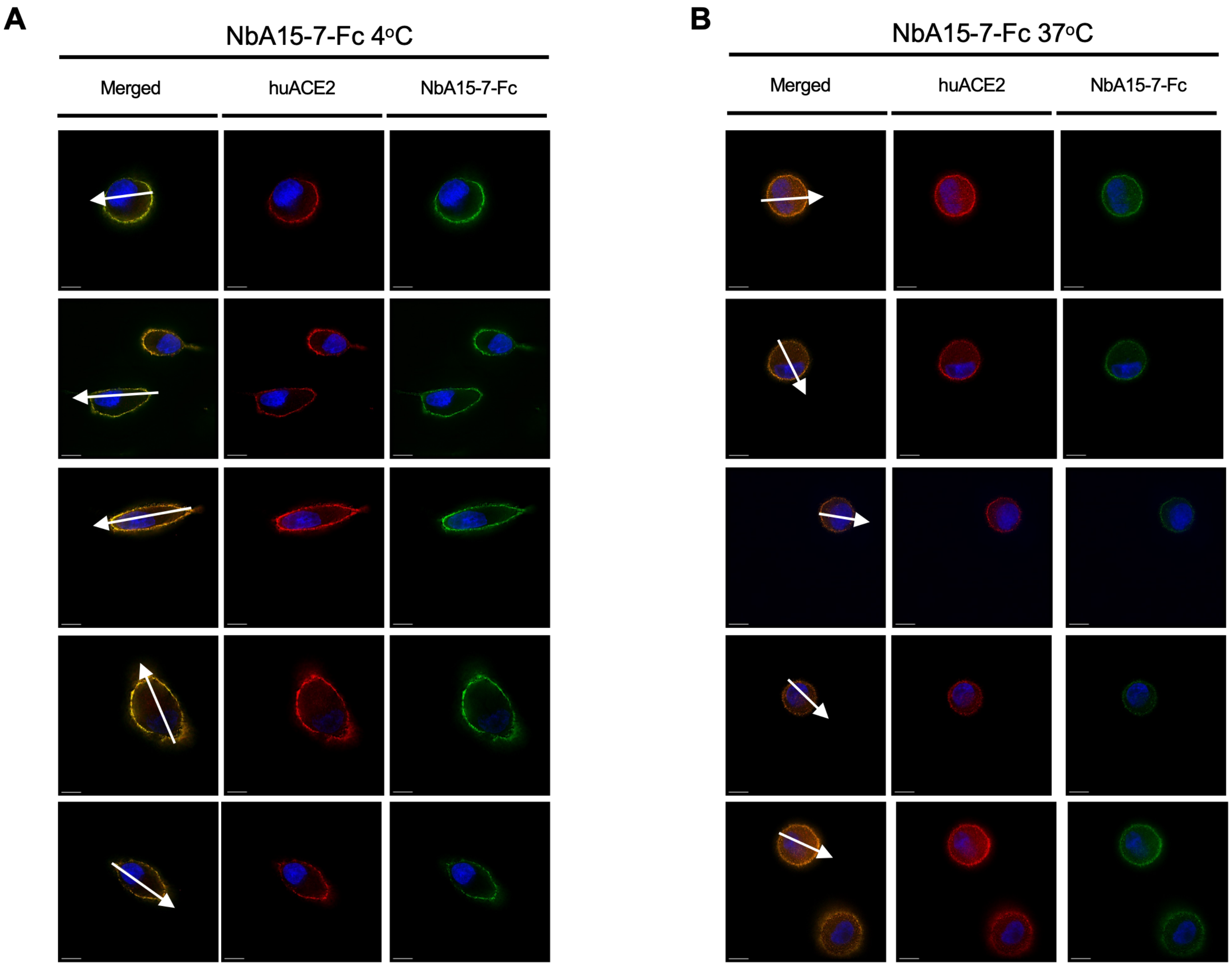

A
NbA15-7-Fc 4°C
Merged
huACE2
NbA15-7-Fc
B
NbA15-7-Fc 37°C
Merged
huACE2
NbA15-7-Fc

**Fig S11: Effect of VHH-Fc fusions on huACE2 localization. (A-B)** Localization of anti-huACE2 VHH-Fc fusion NbA15-7-Fc (Alexa-488, green, right), huACE2-HA (Alexa-594, red, center) and merged (left) following incubation of live A549/hACE2-HA cells at 4$^{o}$C or 37$^{o}$C. Blue stain (DAPI) indicates cell nuclei. Scale bar: 10 μm. Data from a representative experiment (Total experiments=2).

**Fig S12:**

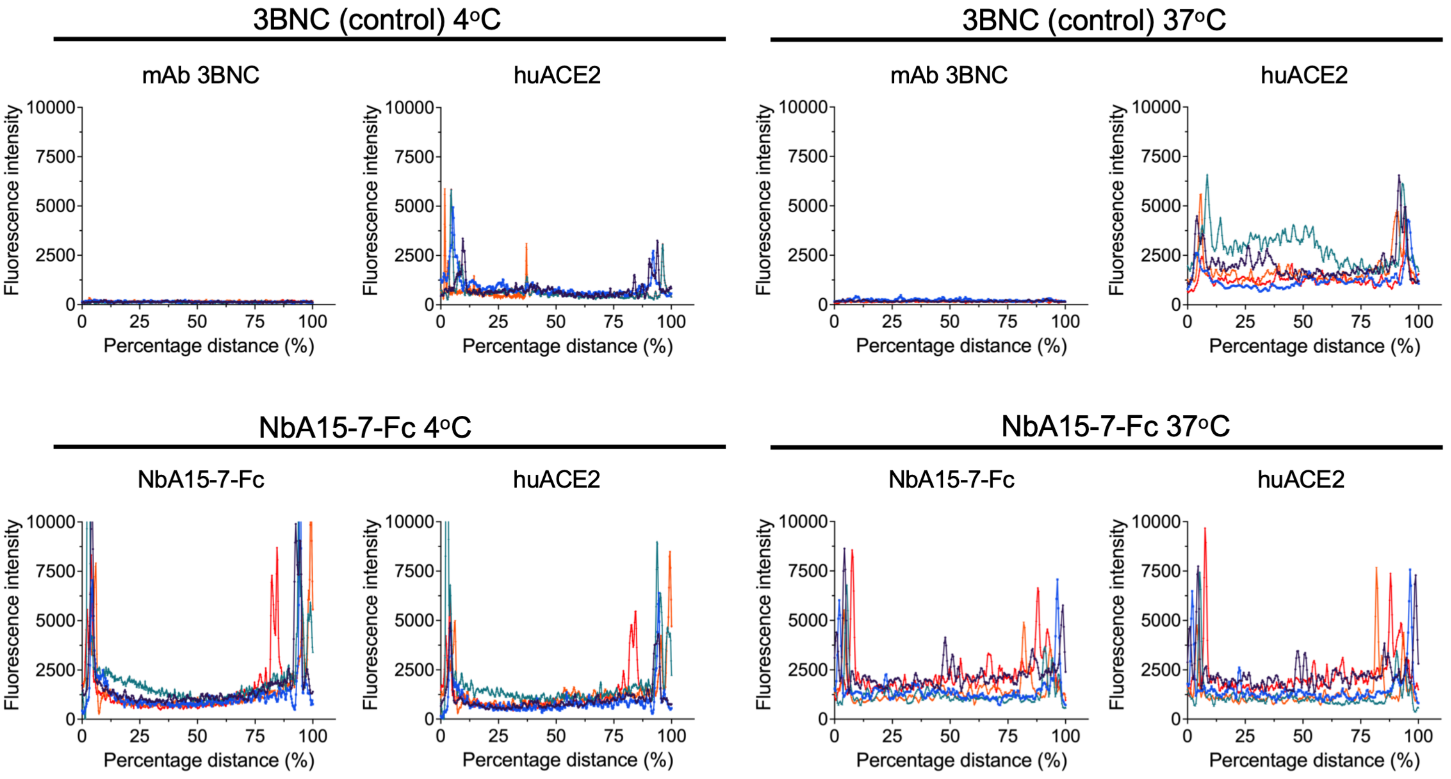
3BNC (control) 4°C
3BNC (control) 37°C
mAb 3BNC
huACE2
NbA15-7-Fc 4°C
NbA15-7-Fc 37°C
NbA15-7-Fc
Fluorescence intensity
Percentage distance (%)
10000
7500
5000
2500
0
25
50
75
100

**Fig S12: Line-scan fluorescence profiles of huACE2 bound to VHH-Fc.**

A549 cells expressing N-term HA tagged huACE2 were incubated with NbA15-7-Fc or the control 3BNC117 (1 µg/ml) for 1h at 4 °C or 37 °C, then fixed, permeabilized, and stained with anti-human IgG-Alexa Fluor 488 (to detect bound VHH-Fc/control mAb) and mouse anti-HA followed by anti-mouse IgG-Alexa Fluor 594 (to detect huACE2). Single z-slices from five representative cells per condition were selected, and membrane-to-membrane line profiles were extracted for each channel. Each trace represents one cell (n = 5). The x-axis shows position along the line scan normalized to cell width (0-100%, membrane to membrane); the y-axis shows fluorescence intensity in arbitrary units (in Fiji). Peaks near 0% and 100% correspond to the plasma membrane, whereas signal across the cell interior indicates intracellular redistribution.

**Fig S13.**

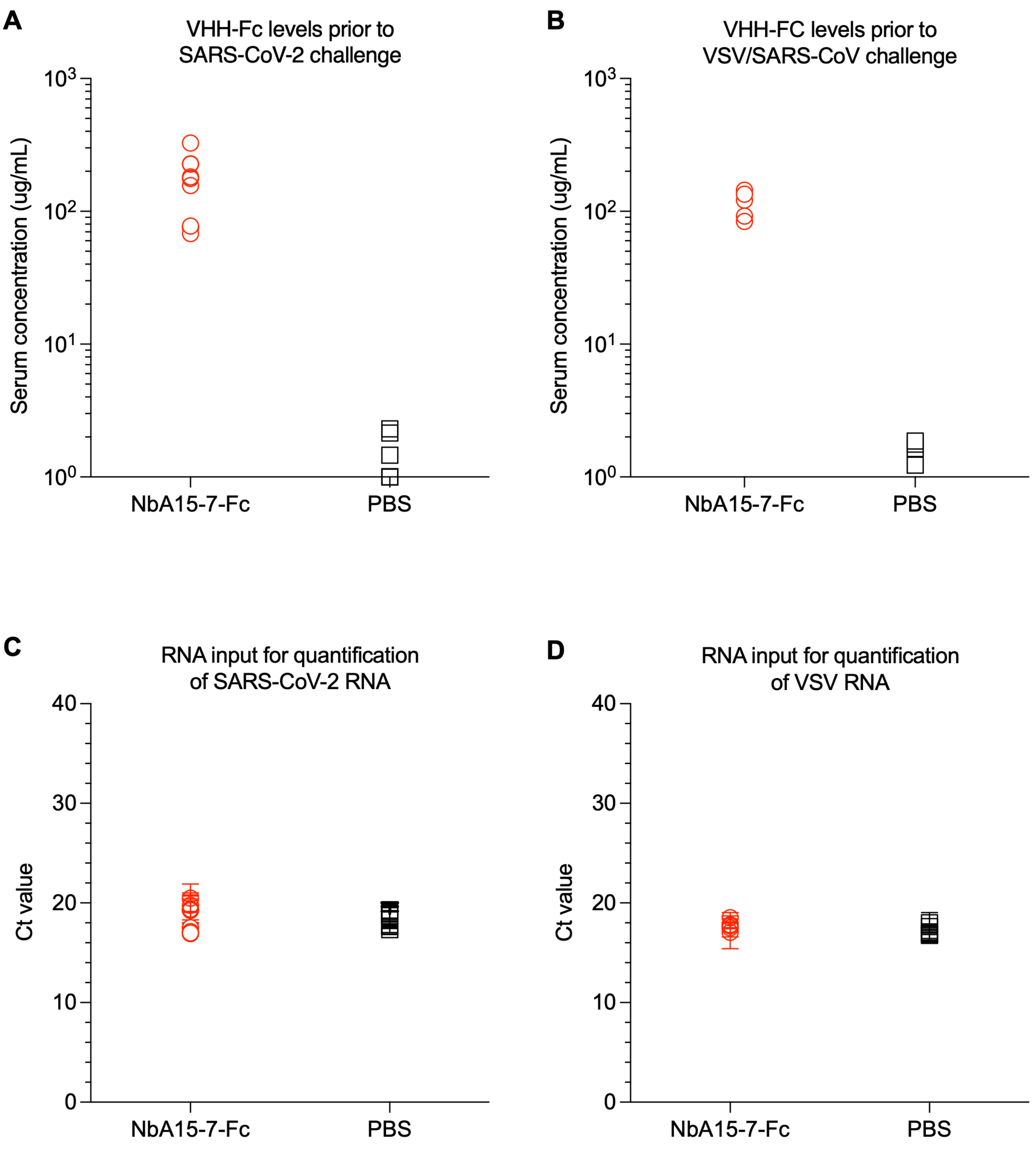
A
VHH-Fc levels prior to SARS-CoV-2 challenge
Serum concentration (ug/mL)
$10^3$
$10^2$
$10^1$
$10^0$
NbA15-7-Fc
PBS
B
VHH-FC levels prior to VSV/SARS-CoV challenge
Serum concentration (ug/mL)
$10^3$
$10^2$
$10^1$
$10^0$
NbA15-7-Fc
PBS
C
RNA input for quantification of SARS-CoV-2 RNA
Ct value
40
30
20
10
0
NbA15-7-Fc
PBS
D
RNA input for quantification of VSV RNA
Ct value
40
30
20
10
0
NbA15-7-Fc
PBS

**Fig S13: *In vivo* pharmacokinetics and therapeutic efficacy of NbA15-7-Fc against SARS-CoV-2 and chimeric VSV/SARS-CoV challenge. (A)** Serum levels of NbA15-7-Fc following i.p. administration in huACE2-KI mice. Mice received either NbA15-7-Fc (red circles) or PBS vehicle control (black squares) prior to SARS-CoV-2 challenge. Sera were harvested from each mouse immediately prior to virus inoculation. Relative viral inhibition curves for sera and standards enabled extrapolation of NbA15-7-Fc serum concentration in each mouse. **(B)** Serum levels of NbA15-7-Fc following i.p. administration in K18-huACE2-IFNAR$^{-}$/$^{-}$ mice. Mice received either NbA15-7-Fc (red circles) or PBS (black squares) prior to challenge with a chimeric VSV/SARS-CoV. Relative virus inhibition of sera samples and standards enabled extrapolation of NbA15-7-Fc serum concentration in each mouse. **(C)** Mean and standard deviation values of qPCR Ct values for GAPDH RNA control from each lung sample showing comparable RNA input during quantification of viral SARS-CoV-2 RNA levels from NbA15-7-Fc-treated (red circles, n=10) and PBS control (black squares, n=10) mice. **(D)** Mean and standard deviation values of qPCR Ct values for GAPDH RNA control from each lung sample showing comparable RNA input during quantification of viral VSV/SARS-CoV RNA levels from NbA15-7-Fc-treated (red circles, n=5) and PBS control (black squares, n=5) mice.
